# Dissociable Multi-scale Patterns of Development in Personalized Brain Networks

**DOI:** 10.1101/2021.07.07.451458

**Authors:** Adam R. Pines, Bart Larsen, Zaixu Cui, Valerie J. Sydnor, Maxwell A. Bertolero, Azeez Adebimpe, Aaron F. Alexander-Bloch, Christos Davatzikos, Damien A. Fair, Ruben C. Gur, Raquel E. Gur, Hongming Li, Michael P. Milham, Tyler M. Moore, Kristin Murtha, Linden Parkes, Sharon L. Thompson-Schill, Sheila Shanmugan, Russell T. Shinohara, Sarah M. Weinstein, Danielle S. Bassett, Yong Fan, Theodore D. Satterthwaite

## Abstract

The brain is organized into networks at multiple resolutions, or scales, yet studies of functional network development typically focus on a single scale. Here, we derived personalized functional networks across 29 scales in a large sample of youths (n=693, ages 8-23 years) to identify multi-scale patterns of network re-organization related to neurocognitive development. We found that developmental shifts in inter-network coupling systematically adhered to and strengthened a functional hierarchy of cortical organization. Furthermore, we observed that scale-dependent effects were present in lower-order, unimodal networks, but not higher-order, transmodal networks. Finally, we found that network maturation had clear behavioral relevance: the development of coupling in unimodal and transmodal networks dissociably mediated the emergence of executive function. These results delineate maturation of multi-scale brain networks, which varies according to a functional hierarchy and impacts cognitive development.

## INTRODUCTION

Executive function (EF) is a broad cognitive domain encompassing top-down modulation of behavior and attention (Jurado & Rosselli, 2007; Gur et al., 2012; Miyake et al., 2000). Compared to earlier-developing cognitive domains, such as episodic memory and motor abilities, EF undergoes extensive, protracted development in youth prior to declining later in life (Fjell et al., 2017; Best & Miller, 2010; Simmonds et al., 2017). Deficits in the development of EF are associated with lower academic achievement (Arffa, 2007; Best et al., 2011), risk-taking behaviors (Casey et al., 2008), and most major psychiatric illnesses (Snyder et al., 2019; Shanmugan et al., 2016; Millan et al., 2012). Because EF requires functional coordination among networks of spatially distributed regions (Niendam et al., 2012; Rottschy et al., 2012; Tan et al., 2018; Murphy et al., 2020), the development of EF is increasingly understood to be dependent on the maturation of multiple large-scale functional networks (Baum et al., 2017; Di Martino et al., 2014; Luna et al., 2015). Consequently, understanding functional network development is essential for any account of how EF develops in youth.

In adulthood, variability in functional network properties is parsimoniously explained by a sensorimotor to association axis (Sydnor et al., 2021); this axis captures variance in network architecture, the spatial ordering of networks on the cortex, and diversity in network-supported faculties and behaviors (Margulies et al., 2016). Further, this axis is thought to support hierarchical information propagation between unimodal networks involved in immediate perception to transmodal networks supporting complex cognition (Mesulam et al., 1998; Murphy et al., 2019; Murphy et al., 2018). Complex cognition, including EF, is hypothesized to partially depend on the segregation of networks located at the top of this functional hierarchy from primary somatosensory activity (Buckner & Krienen, 2013; Mesulam et al., 1998). Indeed, prior studies have reported that the default mode network (DMN; situated at the transmodal end of the axis) segregates from other networks during development, which in turn supports the development of EF (Barber et al., 2013; Sherman et al., 2014; Anderson et al., 2011; Owens et al., 2020). Nonetheless, results remain heterogenous, and the degree to which fundamental properties of cortical hierarchy impacts network development remains unclear (Zhong et al., 2014; Marek et al., 2015; Reinenberg et al., 2015; Dwyer et al., 2014). This lack of consensus across existing work may arise due to two limitations that are shared across prior studies.

First, nearly all studies of functional network development only examine a single network resolution, or scale. Typically, investigators use standard network atlases that specify a single number of functional networks (e.g., 7, 14, or 17). However, it is increasingly recognized that the brain is a multi-scale system, and that studies of a specific resolution of sub-networks may be limited (Brittin et al., 2021; Betzel and Bassett, 2017; Douglas and Martin, 2012; Eickhoff et al., 2018; Breakspear and Stam, 2005). Rather, evidence suggests that brain network organization emerges from neural coordination across overlapping spatial scales (Yeo et al., 2014; Faskowitz et al., 2020; LaBar et al., 1999). Importantly, distinct brain-behavior relationships may be present at different scales (Betzel et al., 2019), with each scale potentially offering complementary information regarding multifaceted processes such as development. As a result, current accounts of brain development that rely on a single network scale are almost certainly incomplete and may hamper our ability to synthesize findings across studies where different scales were examined (de Reus & van den Heuvel, 2013; Arslan et al., 2018).

A second key limitation of prior studies of functional network development is that they have not accounted for individual differences in the spatial layout of brain networks on the cortical mantle. Multiple independent studies in adults using different datasets and distinct methods have provided convergent evidence that there is prominent between-individual variability in the spatial distribution (i.e., the *functional topography*) of large-scale networks on the anatomic cortex (Bijsterbosch et al., 2018; Kong et al., 2019; Li et al., 2019; Gordon et al., 2017). In studies of adults, transmodal association networks tend to have the greatest variability in functional topography (Kong et al., 2019; Li et al., 2019; Gordon et al., 2017; Xu et al., 2016); recent work has shown that this is also true in children and adolescents (Cui et al., 2020). Accounting for such individual variation in functional topography may be critical for understanding the development of coupling between networks, as prior work has shown that differences in topography can be aliased into estimates of connectivity (Bijsterbosch et al., 2018; Burger et al., 2021). Furthermore, individual-specific–or “personalized”–networks may be particularly relevant when evaluating development at multiple scales, as individual variation in topography might depend in part on network resolution (Braga & Buckner, 2017; Steinmetz & Seitz, 1991).

In this study, we sought to understand how multi-scale cortical networks, occupying diverse positions across the sensorimotor-association axis, mature with age to support EF. We evaluated the development of multi-scale personalized networks in a large sample of youth, with the goal of testing three interrelated hypotheses. First, we hypothesized that across scales, patterns of network development would vary across the sensorimotor-association axis, with transmodal networks exhibiting functional segregation relative to unimodal networks. Second, we predicted that transmodal network segregation would in part mediate the maturation of EF in adolescence. Finally, we expected to find evidence of multi-scale network development. Specifically, given the diverse functions supported by brain organization at different scales, we anticipated that different network scales would have distinct associations with both age and EF.

## RESULTS

We studied 693 youths ages 8-23 years from the Philadelphia Neurodevelopmental Cohort, who completed functional MRI (fMRI) at 3T and had 27 minutes of high-quality data (Satterthwaite et al., 2014; Cui et al, 2020). To derive multi-scale personalized functional networks, we used a specialized adaptation of non-negative matrix factorization (NMF) that incorporates spatial regularization (see Methods, **Figure S1**; Lee and Seung, 1999, Li et al., 2017). To ensure correspondence of personalized networks across participants, this process was initialized by creating a group atlas, which was then adapted to each individual’s data (see Methods). To evaluate multiple resolutions, group atlases that included between 2 and 30 networks were created (**Figure 1** and **Figure S2A**). Across this range of scales, reconstruction error declined smoothly (**Figure S2B**).

**Figure 1:**
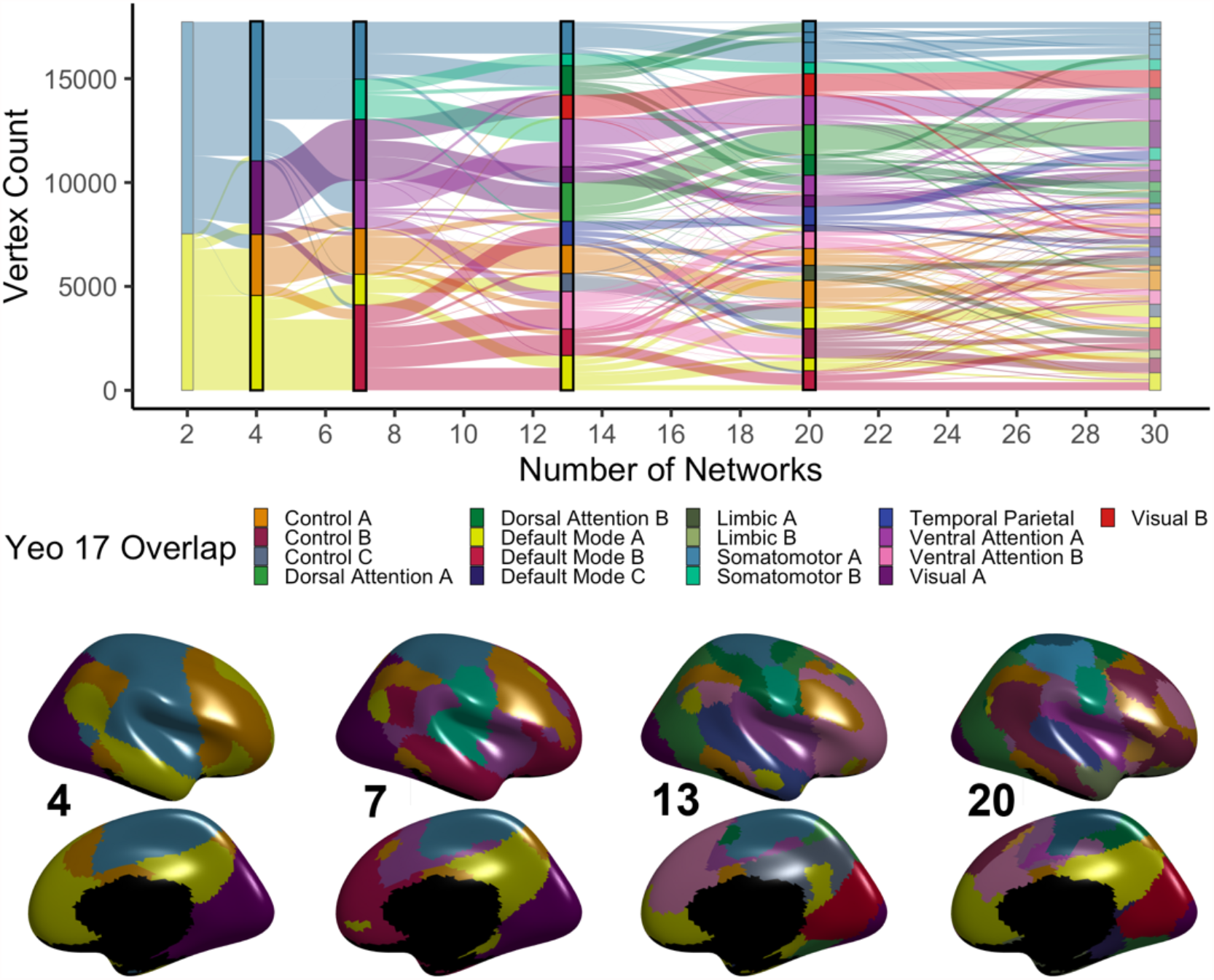
Group consensus functional networks at multiple scales. We used regularized non-negative matrix factorization (see **Supplementary Figure 1**) to derive personalized functional networks at 29 scales (2-30 networks). Tracking network membership of each vertex across scales reveals a nested structure where finer-grained networks gradually emerge from coarse networks (top). Scales 4, 7, 13, and 20 are chosen for visualization; see bottom panel for cortical projections. Colors reflect each network’s predominant overlap with a canonical atlas of 17 functional networks (Yeo et al., 2011).

Examination of multi-scale personalized brain networks revealed prominent differences in person-specific functional neuroanatomy at all scales (**Figure 2A** and **Figure S3**). Prior work at a single scale found that variability in functional neuroanatomy disproportionately localizes to transmodal association cortices (Kong et al., 2019, Cui et al., 2020). Here, to quantify individual variability in network topography, we calculated the median absolute deviation (MAD) of network loadings at each cortical vertex across participants. To verify that variability was consistently greater within transmodal cortex at multiple scales, we compared network MAD at each scale to a widely used map summarizing a unimodal sensorimotor to transmodal association axis of cortical organization, derived from the principal gradient of functional connectivity (Margulies et al., 2016). Using a conservative spin-based spatial randomization procedure that accounts for spatial auto-correlation (Alexander-Bloch et al., 2018), we found that MAD was positively correlated with the unimodal-to-transmodal axis in 27 of the 29 scales evaluated (**Figure 2B**; green). Furthermore, we found that topographic variability became increasingly correlated with this axis at finer scales (**Figure 2C**; *r* = 0.56, *p*_*boot*_ < 0.001). These results demonstrate that variability in functional neuroanatomy is particularly prominent within transmodal cortices at finer-grained network resolutions.

**Figure 2:**
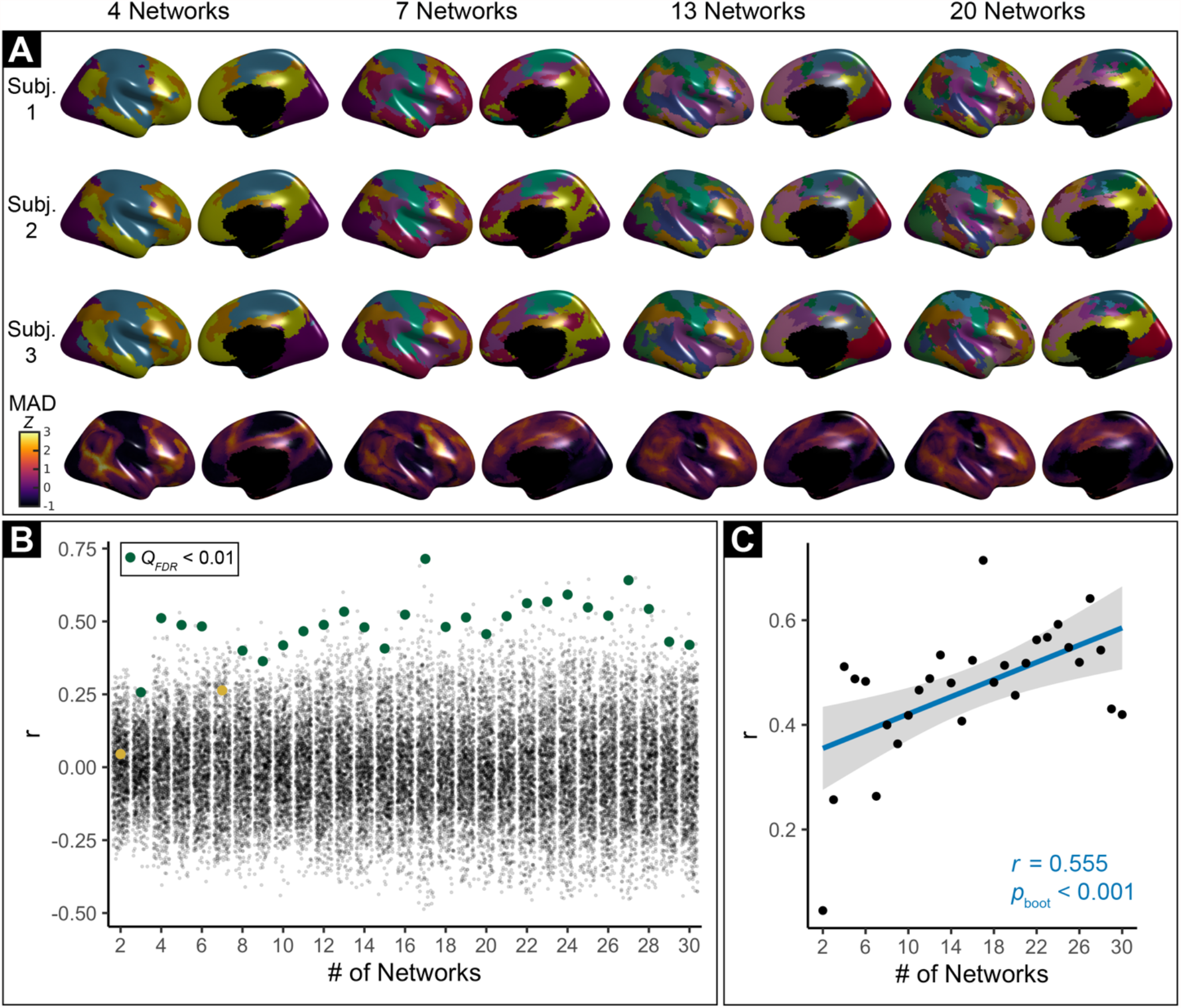
Variability in personalized networks across scales. **A)** Variability in personalized networks is greatest in transmodal cortex across scales. Exemplar personalized networks at scales 4, 7, 13, and 20 are shown for three participants. Prominent individual differences in functional topography are present at all scales, as quantified by median absolute deviation (MAD) of functional network loadings across participants (bottom row, z-scored within each scale). **B)** Variability of functional topography aligns with a unimodal-to-transmodal axis. Spin-tests of the correlation between topographic variability and the principal functional connectivity gradient (Margulies et al., 2016) at each scale reveal that variability is significantly correlated with a unimodal-to-transmodal axis at most scales (green dots = significant correlations; yellow dots = non-significant correlations; black dots = spin-test null correlations). **C)** Greater alignment between a unimodal-to-transmodal axis and topographic variability is present at finer scales. Scatterplot depicts second-order correlation of variability (MAD) and the principal gradient (from **B**) across scales.

### Brain network coupling develops according to a hierarchical unimodal-transmodal axis

Having defined multi-scale personalized networks in a large sample of youth, we next sought to examine how network coupling evolves with age. To summarize functional coupling of each network to other networks, we averaged between-network connectivity values across all personalized networks at each scale **(Figure S4)**. Given that the hierarchical sensorimotor to association axis represents a principal mode of functional coupling in adults (Margulies et al., 2016), but not in infants (Larivière et al., 2020) or children (Dong et al., 2020; Nenning et al., 2020), we hypothesized that age-related changes in between-network coupling would vary according to this axis. Specifically, we expected that functional network development should differ across the axis in a manner that differentiates coupling patterns in unimodal sensorimotor and transmodal association cortices. To test this hypothesis, we first evaluated each networks’ position along this principal axis as the “transmodality” of that network, where higher values correspond to regions located in transmodal association cortices and lower values are assigned to regions in unimodal sensorimotor cortices (**Figure 3A**). Transmodality was operationalized by extracting the average value of a published map of the principal gradient of functional connectivity (Margulies et al., 2016) within each network’s boundaries. We related all network-level age effects to this measure of transmodality.

**Figure 3:**
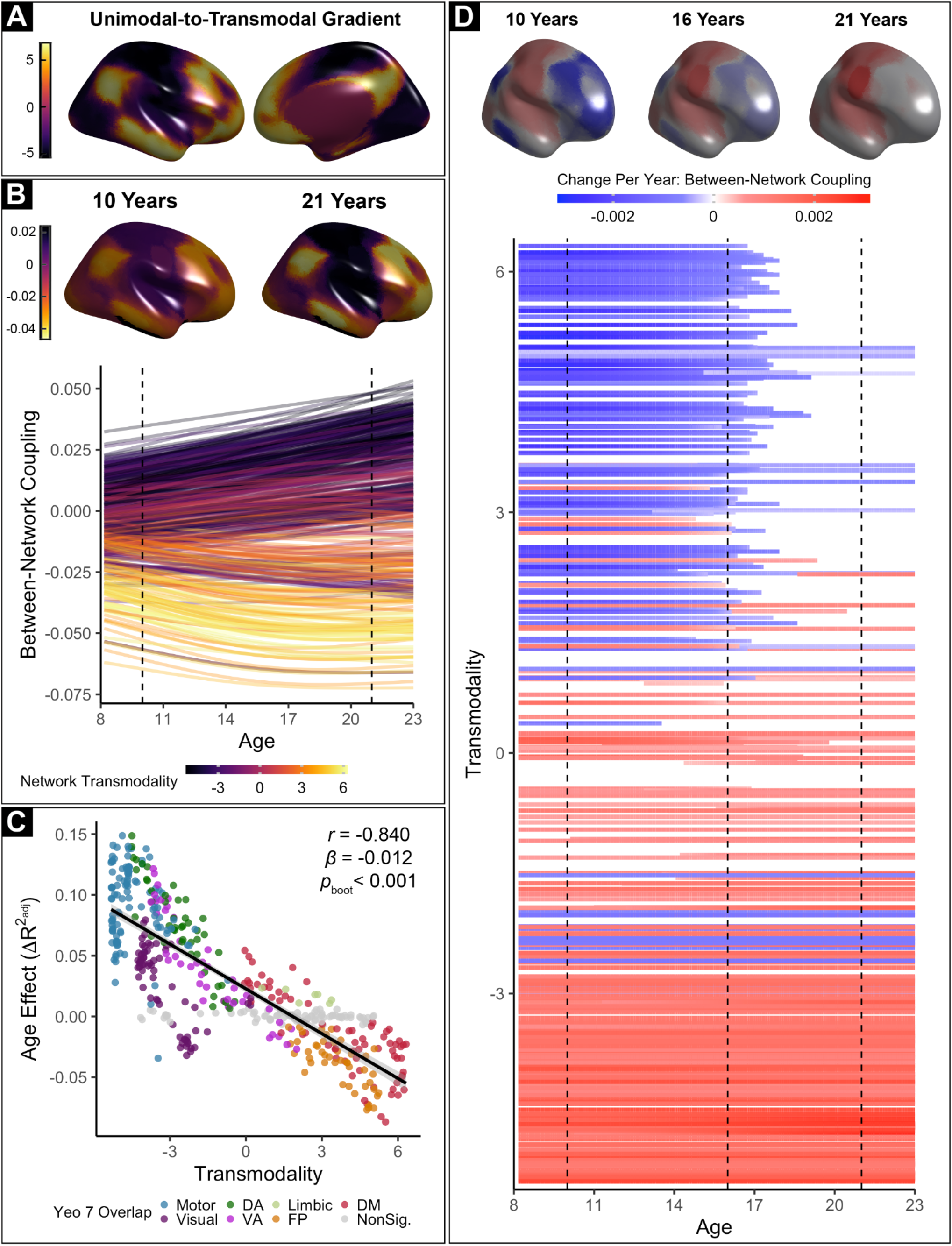
Network development in youth unfolds along a functional hierarchy. **A)** We define functional hierarchy according to the widely used principal gradient of functional connectivity from Margulies et al. (2016), which describes each location on the cortex on a unimodal-to-transmodal continuum (referred to as the “transmodality” value). **B)** Between-network coupling is modeled for every network at each scale using Generalized Additive Models (GAMs) with penalized splines to account for linear and nonlinear effects of age. Each solid line represents the developmental pattern of one network at one scale; colors indicate the average transmodality value within that network. Dashed lines and corresponding brain maps represent estimated between-network coupling at each age, averaged across scales. Between-network coupling of unimodal networks (purple lines) increases with age, indicating increased integration. In contrast, the coupling of transmodal networks (yellow lines) declines with age, reflecting increased segregation. **C)** Age effects of each network (from **B**) are plotted versus their average transmodality (from **A**). Networks that do not display significant change over development are shaded in gray (*Q*_FDR_ > 0.05). The position of each network on the functional hierarchy explains the majority of variance in age effects (*r* = -0.840, *β* = -0.012, *p*_boot_ < 0.001). **D)** We quantified the duration, magnitude, and direction of maturational changes in coupling for each network using the derivatives of the fitted splines (from **B**). Top: annualized change in between-network coupling at 10, 16, and 21 years old, averaged across scales. Bottom: change per year in average between-network coupling of each network across the age range studied; as in **B**, each line represents the developmental pattern of a given network at a single scale. While integration of unimodal networks increases over the entire age range sampled, segregation of transmodal networks generally plateaus near the end of adolescence.

Across all participants and independent of age, average between-network coupling was positive in more unimodal cortices and negative in more transmodal cortices (**Figure 3B**). To rigorously model linear and nonlinear changes in coupling over development, we used generalized additive models (GAMs) with penalized splines to examine how between-network coupling of each network was associated with age. In these models, sex and in-scanner motion were also included as covariates. We found that age-related changes in between-network coupling were largely explained by a network’s position in the functional hierarchy. Between-network coupling of unimodal cortex became more positive at older ages, indicative of greater network integration. In contrast, between-network coupling in association cortex became more negative, reflecting increasing segregation. A network’s position on the functional hierarchy explained most of the variance in observed developmental effects (**Figure 3C**; *r* = -0.84, *p*_boot_ < 0.001). Together, these results suggest that the development of between-network coupling in youth is largely described by dissociable processes of segregation and integration across the sensorimotor to association axis.

Next, we sought to identify intervals of significant age-related change in network coupling. To accomplish this, we calculated the confidence interval of the derivative of the developmental curve for each model. We found that age-related changes in unimodal and transmodal functional networks occurred over different developmental periods: between-network coupling increased in unimodal areas over the entire age range studied, whereas decreases in between-network coupling in transmodal areas did not extend beyond adolescence (**Figure 3D**). Consequently, in addition to differences in the sign of developmental changes described above, the temporal span of maturation in network coupling also systematically varied across the principal axis of cortical organization.

To provide a more nuanced understanding of the maturational changes in between-network coupling described above, we next evaluated development of specific connections between networks. As between-network connections can link networks that have a similar position along the principal axis (i.e., two transmodal association networks) or may alternatively link a unimodal and transmodal network, we calculated the difference of the transmodality values of the two networks connected by each edge. As the principal axis captures variance in cortical coupling, we expected networks similarly positioned along this axis to share a degree of this variance. As expected, we found that networks with similar transmodality values had greater mean coupling, and networks with high transmodality differences tended to have weaker coupling across participants (*r* = -0.57, *p*_boot_ < 0.001; **Figure 4A**). Critically, we additionally found that age-related changes in network edges were also explained by differences in network transmodality (*r* = -0.49, *p*_boot_ < 0.001; **Figure 4B**). Specifically, unimodal-to-unimodal edges tended to strengthen with age, whereas edges that linked unimodal and transmodal networks weakened (**Figure 4C**; *p*_boot_ < 0.001); developmental strengthening of transmodal-to-transmodal edges was present but less prominent. These results demonstrate that functional network development is characterized by increases in coupling between hierarchically similar networks and decreases in coupling between dissimilar networks — yielding increased differentiation along the functional hierarchy with development.

**Figure 4:**
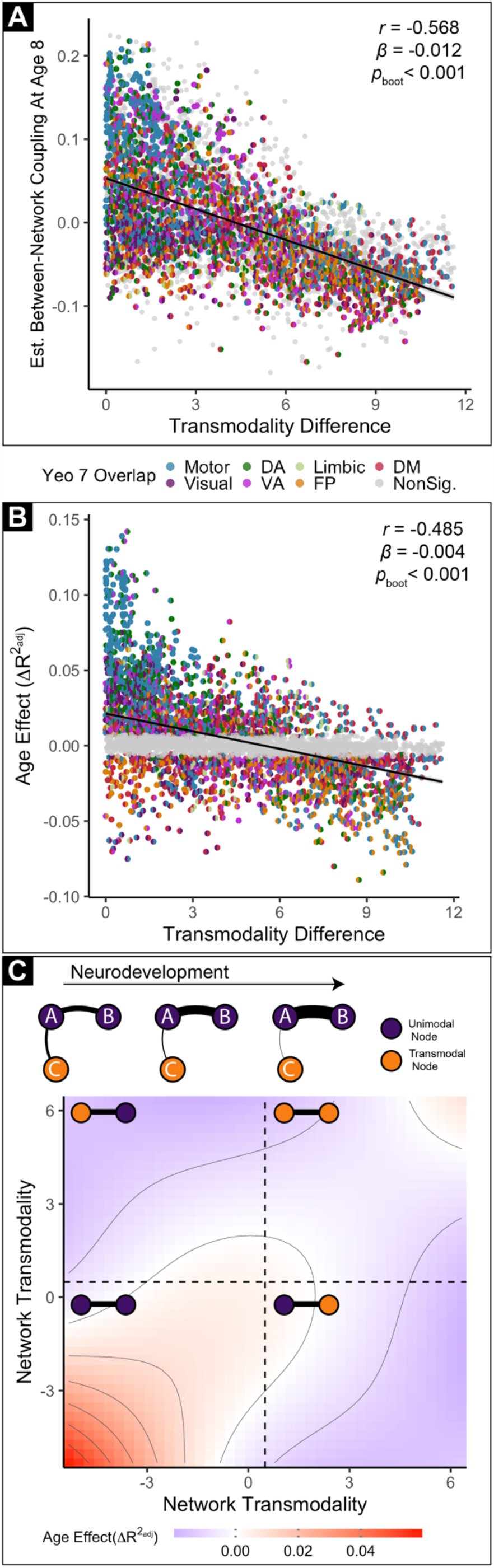
Maturation of between-network coupling aligns with the position of each network in the functional hierarchy. **A)** Mean between-network coupling is largely captured by relative position along the sensorimotor to association axis. The inter-network coupling of each pair of networks at each scale is modeled using a GAM to estimate their values at age 8. Here, those values are plotted versus the difference in the mean transmodality values of the two networks being connected. Each data point represents the coupling of a network pair at a given scale. Each half of the circle is colored according to constituent networks’ maximum overlap with the 7-network solution defined by Yeo et al. (2011); network pairs that do not significantly change with age after FDR correction (*Q* < 0.05) are shaded in gray. As expected, networks at a similar position along the unimodal-to-transmodal axis tend to have higher coupling (*r* = -0.568, *β* = -0.012, *p*_boot_ < 0.001). **B)** Age effects quantifying the development of between-network coupling is similarly aligned with the relative position of networks along the unimodal to transmodal gradient. Age effects of every network pair at each scale are plotted versus their transmodality difference and colored as in **A**. Network pairs without significant age effects are plotted in gray. Developmental effects on pairwise coupling between networks are associated with the transmodality difference between networks (*r* = - 0.49, *p*_boot_ < 0.001). **C)** Top: schematic summarizing developmental effects. Development is associated with strengthening of unimodal-to-unimodal network coupling and weakening of unimodal-to-transmodal coupling; thicker lines represent greater functional coupling. Bottom: topographical plot of the observed age effect as a function of absolute (rather than relative) network transmodality values across all network pairs. Increased coupling with age between functionally similar networks is prominent for unimodal networks (bottom left), and less prominent for transmodal networks (top right). Age-related decreases in coupling occur in unimodal-transmodal network pairs (top left and bottom right).

It should be noted that previous studies have documented that the physical distance between two brain regions explains the patterning of functional maturation across network edges (Fair et al., 2007, Power et al., 2010; Ma et al., 2021). As the principal axis is related to the intrinsic geometry of the cortex (Oligschläger et al., 2017; Huntenburg et al., 2018), we sought to verify that the effects of transmodality difference described above were not better explained by physical distance. To do so, we compared the correlation between age effects and Euclidean distance with the relationship between age effects and transmodality difference. While the correlation between Euclidean distance and age effects was significant (*r* = -0.11, *p*_boot_ < 0.001; **Figure S5**), it was substantially weaker than that observed for transmodality difference (*r* = - 0.49, *p*_boot_ < 0.001) and the effect of transmodality difference remained significant while co-varying for Euclidean distance (partial *r* = -0.45, *p* < 0.001). This result suggests that although the physical distance spanned by a functional connection is weakly related to its developmental pattern, developmental effects are better explained by the functional distance that an connection spans across the sensorimotor to association axis.

### Development has dissociable signatures at different networks and scales

The above results demonstrate that functional network development is largely captured by a network’s position on a hierarchical axis of unimodal-to-transmodal function. However, these analyses are agnostic to the multi-scale nature of the personalized brain networks that we constructed. As a next step, we evaluated whether developmental effects were dependent on network scale. Initial inspection revealed that the relationship between age and between-network coupling varied systematically as a function of scale, with greater age effects in the somatomotor cortex at finer network scales (**Figure 5A**). To quantify scale effects while controlling for within-subject correlations over scales, we used generalized estimating equations (GEEs) with exchangeable correlation structures at each cortical vertex. We found that the effect of scale on between-network coupling was strongest in the somatomotor cortex (**Figure 5B**). Furthermore, we found evidence that scale moderated age effects, with maximal scale-by-age interactions being observed in the somatomotor cortex (**Figure 5C**).

**Figure 5:**
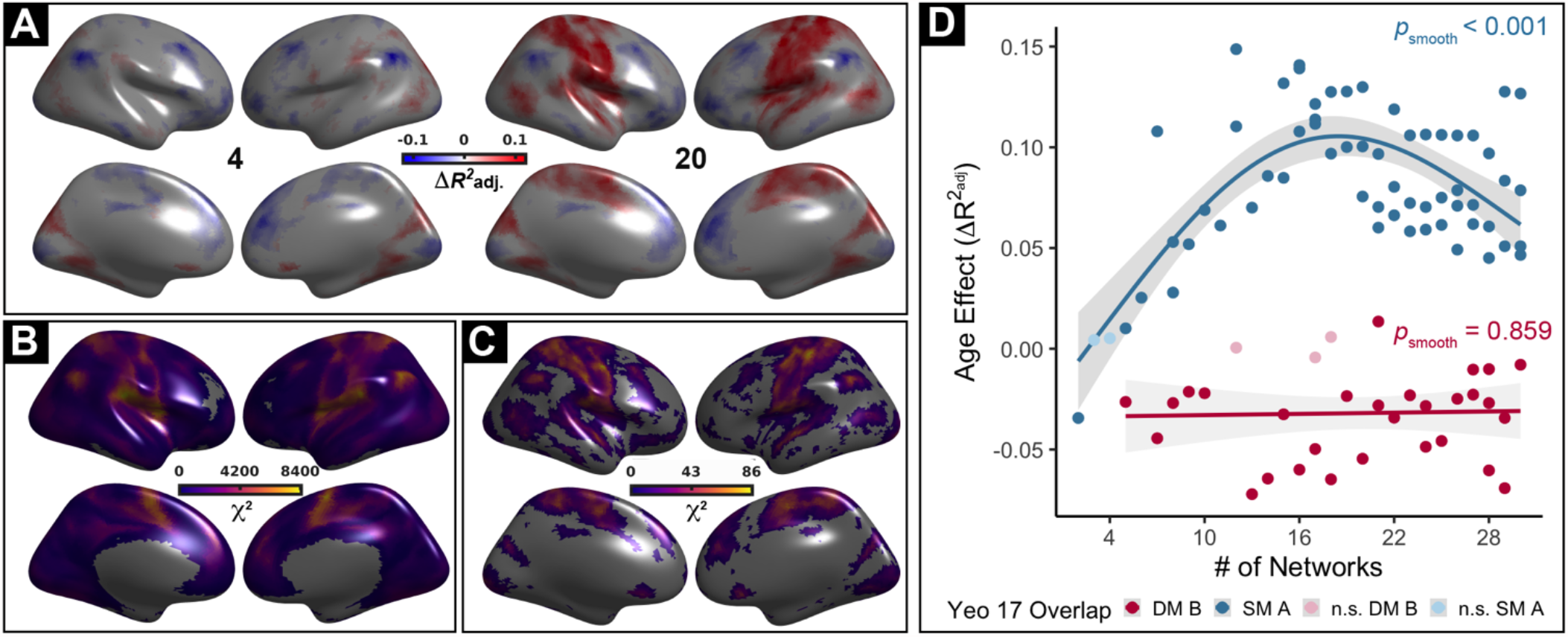
Impact of network scale on development is maximal in somatomotor cortex. **A)** The effect of age on average vertex-wise between-network coupling at two scales (4 and 20). Age effects are modeled using GAMs with penalized splines; thresholded at *Q*_FDR_ < 0.05. Scale-dependent age effects can be observed in somatomotor cortex: while no developmental increase in between-network coupling was seen in somatomotor cortex at scale 4, such an increase is evident at scale 20. **B)** Across ages, between-network coupling of the somatomotor cortex is strongly influenced by scale. Generalized estimating equations (GEEs) reveal that the effect of scale (*χ*^2^) differentially influences the strength of between-network coupling across the cortex. Locations within unimodal somatomotor cortex exhibit the strongest scale-dependence in their mean between-network coupling (*Q*_FDR_ < 0.05). **C)** Scale differentially impacts age-dependent developmental associations with coupling across the cortex. GEEs are used to examine the degree of scale-moderated developmental effects (age-by-scale interaction; thresholded at *Q* < 0.05); maximal effects are present in the somatomotor cortex. **D)** Scale differentially impacts age-dependent developmental effects in transmodal and unimodal networks. Specifically, age effects in unimodal somatomotor networks tend to be more scale-dependent than those in transmodal networks. The effect of age across scales is plotted for networks predominantly overlapping with the most unimodal (blue; Somatomotor A) and most transmodal (red; Default Mode B) networks.

To further understand these scale-dependent age effects, we compared the age effect across scales for networks that fall at opposite ends of the sensorimotor-to-association axis. Specifically, at each scale we identified networks that aligned most closely with the somatomotor-A network and the default mode-B network from the commonly used atlas defined by Yeo et al. (**Figure 5D**). This comparison revealed that age effects within the somatomotor network were highly scale-dependent, with greater increases in between-network coupling with age at finer scales. In contrast, default mode networks demonstrated consistent developmental segregation across scales. These results suggest that age-related changes in network coupling are differentially impacted by scale across the principal axis.

### Multi-scale network coupling is associated with executive function

Having delineated developmental changes in between-network coupling, we next sought to understand the implications for individual differences in executive function (EF). First, we modeled the association between network coupling and EF, controlling for developmental effects by including age as a penalized spline; other model covariates included sex and motion as in prior analyses. We found that the relationship between EF and between-network coupling was quadratically related to transmodality (**Figure 6A**; *p*_boot_ = 0.003); this quadratic pattern was markedly different than the linear relationship between transmodality and age effects (see **Figure 3C** for comparison). Specifically, decreased between-network coupling at both extremes of the principal axis was associated with greater EF, with maximal effects being seen in somatomotor and default mode networks. In contrast, greater coupling of several visual, ventral attention, and fronto-parietal networks were associated with greater EF.

**Figure 6:**
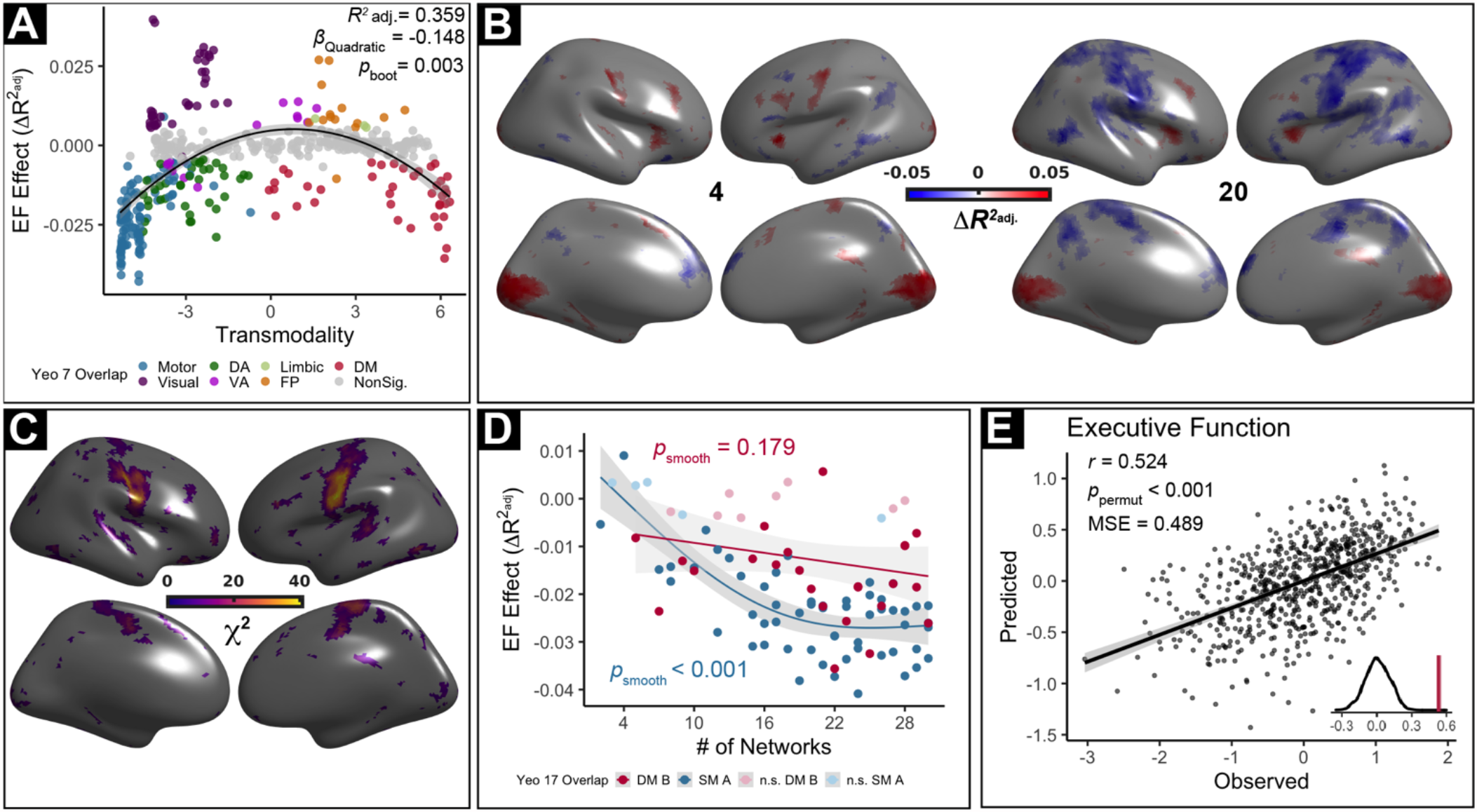
Multi-scale network coupling is associated with executive function. **A)** Network-level relationships between coupling and EF are quadratically related to transmodality. Specifically, segregation of both somatomotor and default-mode networks is associated with better EF. These associations with EF are dissociable from normative developmental effects (**Figure 3C**) where default mode segregation and somatomotor integration are observed. **B)** Analyses at scales 4 and 20 reveal differing associations with EF. While between-network coupling of visual, insular, and dorsolateral prefrontal cortical areas is consistently associated with greater EF (*Q*_FDR_ < 0.05), opposite associations with EF were present in motor cortex at coarse and fine scales. **C)** Tests of age-by-scale interactions using GEEs reveal that scale effects are strongest in the somatomotor cortex. **D)** Scale differentially impacts EF associations with coupling in transmodal and unimodal networks. As for age, effects in unimodal somatomotor networks tend to be more scale-dependent than those in transmodal networks. The effect of age across scales is plotted for networks predominantly overlapping with the most unimodal (blue; Somatomotor A) and most transmodal (red; Default Mode B) of the Yeo 17 networks. **E)** Complex patterns of multi-scale coupling between personalized networks accurately predicts EF in unseen data. Cross-validated ridge regression with nested parameter tuning was used to predict EF of unseen data using each participant’s multivariate pattern of coupling across scales.

To further understand these effects, we next performed high-resolution analyses at each cortical vertex to better understand associations between EF and between-network coupling across scales. Consistent with network-level results, reduced between-network coupling in default mode regions like the medial prefrontal cortex and precuneus was associated with greater EF across scales (**Figure 6B**). In contrast, greater between-network coupling in the dorsolateral prefrontal cortex, anterior insula, and calcarine fissure were associated with greater EF across scales. Somatomotor cortices again exhibited scale-dependent associations: higher between-network coupling in somatomotor cortex was associated with reduced EF, but only at finer scales. To further assess the impact of network scale, we used GEEs to examine whether there was an interaction between EF and scale on between-network coupling at each cortical location. This analysis revealed prominent scale effects, primarily in somatomotor cortices (**Figure 6C**). To further illustrate the differential effects of network scale, we again contrasted networks that lie at opposite ends of the unimodal-transmodal axis (**Figure 6D**). We found that network scale did not moderate the association between default mode network coupling and EF; greater default mode segregation was associated with better EF across all scales. However, somatomotor network associations with EF were highly dependent on network scale.

Having found evidence of both scale-dependent and scale-independent associations between EF and network coupling, we next examined the degree to which these complex patterns of coupling could jointly predict individual differences in EF. To do so, we fit a multivariate ridge regression model to predict EF using data from all scales, while controlling for age and in-scanner motion. We found that this multivariate model accurately predicted EF in unseen data (see Methods; **Figure 6E**; *r* =0.524, *p*_permut_ < 0.001). These results emphasize that EF is supported by multi-scale patterns of functional coupling.

### Multi-scale network development mediates the development of executive function

The prior analyses revealed associations between coupling and EF while controlling for age. However, as in prior studies, we found EF develops dramatically in youth (**Figure 7A**; *r* = 0.41, *p* < 0.001). Accordingly, we evaluated whether maturation in coupling between multi-scale personalized networks mediated the development of EF with age (**Figure 7B**). Critically, our prior results revealed that association networks become more *segregated* with age, and that greater segregation is associated with *better* EF. In contrast, somatomotor networks become more *integrated* with age, but greater integration is associated with *lower* EF. Consequently, we anticipated that networks at opposite ends of the unimodal-transmodal functional axis would have dissociable mediation effects. Mediation analyses revealed that while decreased coupling in more transmodal networks supports the development of EF, increased age-related coupling in more unimodal networks suppresses the development of EF (**Figure 7C**; *r* = 0.68, *p*_boot_ < 0.001). Finally, we tested whether these mediation effects were scale dependent. We found that scale moderated these mediation effects within unimodal somatomotor networks. At finer scales, unimodal network integration attenuated developmental gains in EF (**Figure 7D**). Together, our findings suggest that multi-scale development of network coupling mediates the development of EF. Notably, the impact of each network’s development on EF was explained by its position on the sensorimotor to association axis.

**Figure 7:**
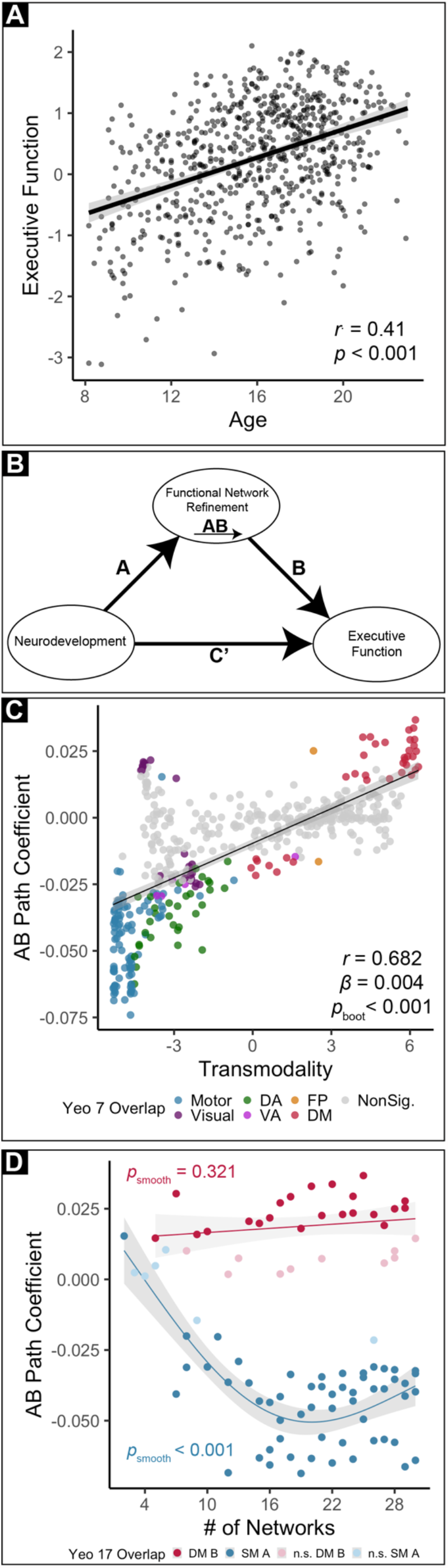
Multi-scale functional maturation mediates the development of executive function in youth. **A)** EF improves with age. As expected, age is significantly correlated with EF in youth (*r* = 0.41, *p* < 0.001). **B)** We evaluated a mediation model in which the association between age and EF is mediated by functional network refinement. **C)** Network-level mediation effects align with functional hierarchy. Negative mediation effects (AB coefficients) are present in unimodal networks, whereas positive mediation effects are present in transmodal networks. This pattern suggests that greater integration of unimodal networks with age is associated with lower EF, whereas greater segregation of transmodal networks is associated with higher EF. **D)** Mediation effects differ by scale in unimodal networks. Mediation weights are plotted as a function of scale for networks predominantly overlapping with the most unimodal and transmodal networks. Instances where network mediation effects do not survive FDR correction are denoted by desaturated points. Again, the strength of mediation effects is scale dependent in somatomotor networks. In somatomotor networks, the strength of negative mediation effects is greatest at finer scales.

## DISCUSSION

In this study, we demonstrated that variation in the development of person-specific functional networks is intrinsically related to fundamental properties of brain organization. Specifically, we found that developmental patterns differentially unfold along the hierarchical sensorimotor to association axis of organization: unimodal somatomotor networks became more integrated with age, while transmodal association networks became more segregated. This dissociable pattern of maturation had unique relevance for the development of cognition: while greater segregation of association networks was associated with better EF, developmental integration of unimodal networks was associated with worse EF. By examining functional network development and associations with EF across a range of macroscale networks, we additionally identified scale-dependent effects, which were predominantly present in somatomotor networks. Taken together, these results provide a new framework that incorporates multi-scale cortical organization for understanding how functional network maturation allows for the development of cognition in youth.

### Functional network development differs by position in a unimodal to transmodal hierarchy

Previous work in adults (Gordon et al., 2017; Kong et al., 2019; Li et al. 2019) has established that between-individual variability of functional topography is greatest in association cortex. In our prior report (Cui et al., 2020), we demonstrated that this is also true in youth. Such marked variability of functional topography in transmodal cortices may be a result of protracted and environmentally sensitive development in higher-order cortices, facilitating continuous adaptation to individual-specific needs (Buckner & Krienen, 2013; Vainik et al., 2020). Here, we extended prior findings by demonstrating that topographic variability aligns with a unimodal-to-transmodal axis across multiple network scales. Furthermore, we found that variability of functional topography increasingly localizes to transmodal association cortex as the number of functional networks increases. As this scale-dependency might be just one of many shifts in between-participant variability over scales (Bijsterbosch et al., 2020; Betzel et al., 2019), our results highlight the importance of scale and precision functional mapping techniques for investigations of individual differences in functional network coupling.

We found strong evidence that developmental changes in between-network coupling align with sensorimotor to association axis. Even prior to adolescence, unimodal networks tended to have greater between-network coupling, which was primarily driven by their coupling with other unimodal networks. In contrast, transmodal networks were more functionally segregated even among the youngest of our participants. From ages 8 to 23 years, this pattern became more prominent: between-network coupling further strengthened in unimodal networks and weakened with age in transmodal networks. Together, these developmental effects served to further distinguish the functional hierarchy that is now well described in adults, and broadly aligns with recent reports using independent methods and datasets (Dong et al., 2020; Nenning et al., 2020). This functional differentiation of cortical hierarchy over development is consistent with evidence that cortical myeloarchitecture further differentiates between unimodal and transmodal regions during adolescence (Paquola et al., 2019), and that transmodal structural networks become increasingly dissimilar from unimodal networks with age (Park et al., 2021). Coupling between hierarchically similar networks may be partially attributable to the propagation of cortical waves along functional hierarchies (Mitra and Raichle, 2016; Matsui et al., 2016; Gu et al., 2021); however, additional research is needed to examine how such waves evolve in development. Taken together, our results suggest that functional network development in youth both aligns with and strengthens the sensorimotor to association axis seen in adulthood.

### Functional network differentiation supports executive function

EF is supported by coordinated recruitment of distributed networks of brain regions (Shine et al., 2019; Satterthwaite et al., 2013a; Tan et al., 2018; Murphy et al., 2020). We found that *segregation* of networks located at the two opposing ends of the sensorimotor to association axis (i.e., somatomotor and default mode networks) was associated with cognitive performance. Conversely, we demonstrated that increased *integration* of networks more centrally positioned within the axis supported EF. As such, two dissociable patterns of normative network development observed across the cortical functional hierarchy differentially impact the development of EF. Specifically, whereas normative developmental segregation of transmodal association networks was positively associated with EF, unimodal integration was positively associated with age but *negatively* associated with EF. These results in part explain the existing heterogeneous literature, which has reported that refinement of both functional network segregation and integration is important for neurocognitive development (Baum et al., 2017; Sherman et al., 2014; Grayson and Fair, 2017; He et al., 2019). However, our results also specify that the degree to which developmental integration versus segregation is advantageous for EF may largely depend on a network’s role within the functional hierarchy.

That both somatomotor and DMN segregation were associated with greater EF accords with recent work demonstrating that the overall balance of network activity shifts across the functional hierarchy when individuals are engaged in externally oriented versus internally guided cognition. Prior work has shown that localized activity within unimodal networks at the bottom of the hierarchy supports cognition when it is reliant on immediate perceptual input (Murphy et al., 2019). In contrast, greater segregation of unimodal networks from transmodal networks supports cognition that is dependent on internally-oriented processing, including memory or theory of mind (Barber et al., 2013; Murphy et al., 2018; Murphy et al., 2019). Furthermore, the association between EF and integration of control networks situated more centrally in the hierarchy is supported by prior literature emphasizing the role of these networks in top-down control (Niendam et al., 2012; Cole et al., 2013; Marek & Dosenbach, 2018). Speculatively, these results suggest that functional segregation at the extremes of the functional hierarchy, in tandem with integration of control networks situated in the middle of the hierarchy, may serve to reduce cross-modal interference (Sonuga-Barke & Castellanos, 2007; Bomyea et al, 2018) while facilitating coordination of brain networks specialized for top-down cognitive control (Cole et al., 2013; Marek & Dosenbach, 2018).

This heterogeneous impact of functional network maturation on EF is particularly relevant when considered in the context of the broader lifespan. In late life, cortical networks re-integrate, losing the segregation that is achieved earlier in the lifespan (Betzel et al., 2014; Chan et al., 2014, Park et al., 2004). Notably, reduced functional segregation has been shown to mediate cognitive decline in both normal aging and neurodegenerative disease (Goh, 2011; Geerligs et al., 2014; Cassady et al., 2021). In parallel to this literature in aging, we found evidence for integration of networks in youth in fine-grained somatomotor networks, which was similarly associated with reduced cognitive performance. A potential implication of these results is that network re-integration associated with cognitive decline in late life may begin far earlier than previously appreciated in fine-grained, early-maturing somatomotor networks.

### Multi-scale patterns of network development impact executive function

Prior work has primarily investigated organizational regimes of 2 (Doucet et al., 2011), 3 (Margulies et al., 2016), 4 (Shokri-Kojori et al., 2019), 5 (Seeley et al., 2009), 6 (Uddin et al., 2019), 7 (Yeo et al., 2011), 13 (Gordon et al., 2014; Power et al., 2011), and 17 (Yeo et al., 2011) functional subdivisions of the brain. Distinguishing the specific role of scale in brain organization is critical for studies of the developing functional hierarchy, as finer scales systematically capture shorter “neural bridges” (Mesulam, 1998) across the functional hierarchy. In other words, as higher network resolutions distinguish increasingly similar sub-networks, finer scales ultimately capture functional interactions between networks that are more proximate in the functional hierarchy. At the coarsest scale of two functional subdivisions, between-network coupling reflects interactions between only a single unimodal-like and transmodal-like network. At this resolution, network segregation between these two broad classes of cortices increased with age. In contrast, finer scales revealed that along with overall developmental segregation of unimodal and transmodal networks, there is prominent integration of functionally similar, finer-grained networks.

We observed independent effects of scale on both development and cognition across the functional hierarchy. In development, integration of somatomotor networks was most prominent at finer scales. In contrast, no such scale dependence was seen in the most transmodal association networks. The scale-invariance of developmental effects in association networks like the DMN may underlie the relative consistency of reported DMN segregation in prior studies (Fair et al., 2007; Sherman et al., 2014; Satterthwaite et al., 2013a). A divergent pattern of scale-dependence was present in associations with EF: whereas the normative integration of finer-grained somatomotor networks was associated with worse EF in development, normative segregation of transmodal networks across scales facilitated the development of EF. Notably, it would be difficult to observe such distinct effects if network coupling were only considered at a single scale.

### Limitations

Several limitations to the current study should be noted. First, there are undoubtedly individual differences in the pace of brain development (Tooley et al., 2021). Future longitudinal studies will be critical for understanding individual deviations in network maturation and psychopathological consequences (Di Martino et al., 2014). Second, as children tend to move more during MRI scans, in-scanner head motion continues to be a concern for all neuroimaging studies of development (Satterthwaite et al., 2013b). Here, we rigorously followed the best practices for mitigating the influence of head motion on our results, including use of a top-performing preprocessing pipeline and co-varying for motion in all hypothesis testing (Ciric et al., 2018). Use of these conservative procedures limits the possibility that reported findings are attributable to in-scanner motion. Third, we used data combined across three fMRI runs, including two where an fMRI task was regressed from the data (Fair et al, 2007). This choice was motivated by studies that have shown that functional networks are primarily defined by individual-specific rather than task-specific factors and that intrinsic networks during task performance are similarly organized to those at rest (Gratton et al., 2018). Importantly, by including task-regressed data, we were able to generate individualized networks with 27 minutes of high-quality data. Prior work suggests that parcellations created using a timeseries of this length show high concordance with those generated using 380 minutes of data (Laumann et al., 2015). Fourth, we studied multi-scale organization in the spatial domain; the brain also exhibits multi-scale organization in the temporal domain (Palva & Palva, 2018; Buzsaki & Draguhn, 2004; Smith et al., 2012, Vidaurre et al., 2017). Future investigations using tools with greater temporal resolution may be critical for simultaneously describing spatial and temporal multi-scale organization. Finally, the maturation of subcortical structures is a critical component of neurodevelopment (Mills et al., 2014; Sommerville et al., 2009). Recent advances in precision functional mapping of subcortical regions (Greene et al., 2020; Sylvester et al., 2020) present a excellent opportunity to delineate the role of subcortical functional coupling in neurocognitive development.

## Conclusion

We leveraged advances in machine learning to elucidate divergent patterns of functional network development and to establish their relevance for cognition. These results are important for understanding the developmental refinement of cortical hierarchy that is prominent in healthy adults. Moving forward, the process of this refinement may be critically important for understanding executive dysfunction in those affected by mental illness. Examining abnormalities of functional network re-organization in longitudinal clinical samples will provide an important opportunity to test the hypothesis that insufficient maturational segregation of transmodal networks confers risk to diverse psychiatric disorders. Indeed, existing research suggests that abnormalities associated with cross-disorder psychopathology are predominantly present at the transmodal end of the functional hierarchy (Shanmugan et al., 2016; Romer et al., 2020; Parkes et al., 2021), and that diverse psychopathology is associated with attenuated segregation of higher-order networks (Xia et al., 2018). Eventually, understanding the normative development of individualized networks may be a critical prerequisite for guiding personalized neuromodulatory interventions targeting both individual-specific functional neuroanatomy and developmental phases with amenable plasticity.

## ACKNOWLEDGEMENTS

This study was supported by grants from the National Institute of Health: F31 MH123063-01A1 (A.P.), R01MH113550 (T.D.S. & D.S.B.), R01EB022573 (C.D., Y.F., & T.D.S.), R01MH120482 (T.D.S. & M.M.), RF1MH116920 (T.D.S. & D.S.B.), R37MH125829 (D.F. & T.D.S), R01MH112847 (R.T.S. & T.D.S.), and T32MH014654 (B.L.). V.J.S. was supported by a National Science Foundation Graduate Research Fellowship (DGE-1845298). L. P. was supported by the 2020 NARSAD Young Investigator Grant from the Brain & Behavior Research Foundation. The PNC was supported by MH089983 and MH089924. Additional support was provided by the Lifespan Brain Institute at Penn and the Children’s Hospital of Philadelphia and the Dowshen Program for Neuroscience. The content is solely the responsibility of the authors and does not represent the official views of any of the funding agencies.

## AUTHOR CONTRIBUTIONS

Conceptualization, T.D.S. and A.R.P.; Methodology, T.D.S., A.R.P., B.L., R.T.S., and S.M.W.; Software, A.R.P., Z.C., H.L., Y.F., T.D.S., A.A., B.L., and A.F.-A-B.; Validation; Z.C., B.L., and M.A.B.; Formal Analysis, A.R.P.; Resources, R.C.G. and R.E.G., Data Curation, T.D.S. and A.A., Writing - Original Draft, A.R.P., B.L., M.A.B., and T.D.S.; Writing – Review and Editing, A.R.P., V.J.S., T.D.S., Z.C., A.F.A-B., C.D., D.A.F., R.C.G., R.E.G., H.L., M.P.M., T.M.M., K.M., L.P., S.L.T-S., S.S., R.T.S., S.M.W., D.S.B., Y.F., and T.D.S.; Visualization, A.R.P.; Supervision, T.D.S.

---

**Figure S1:**
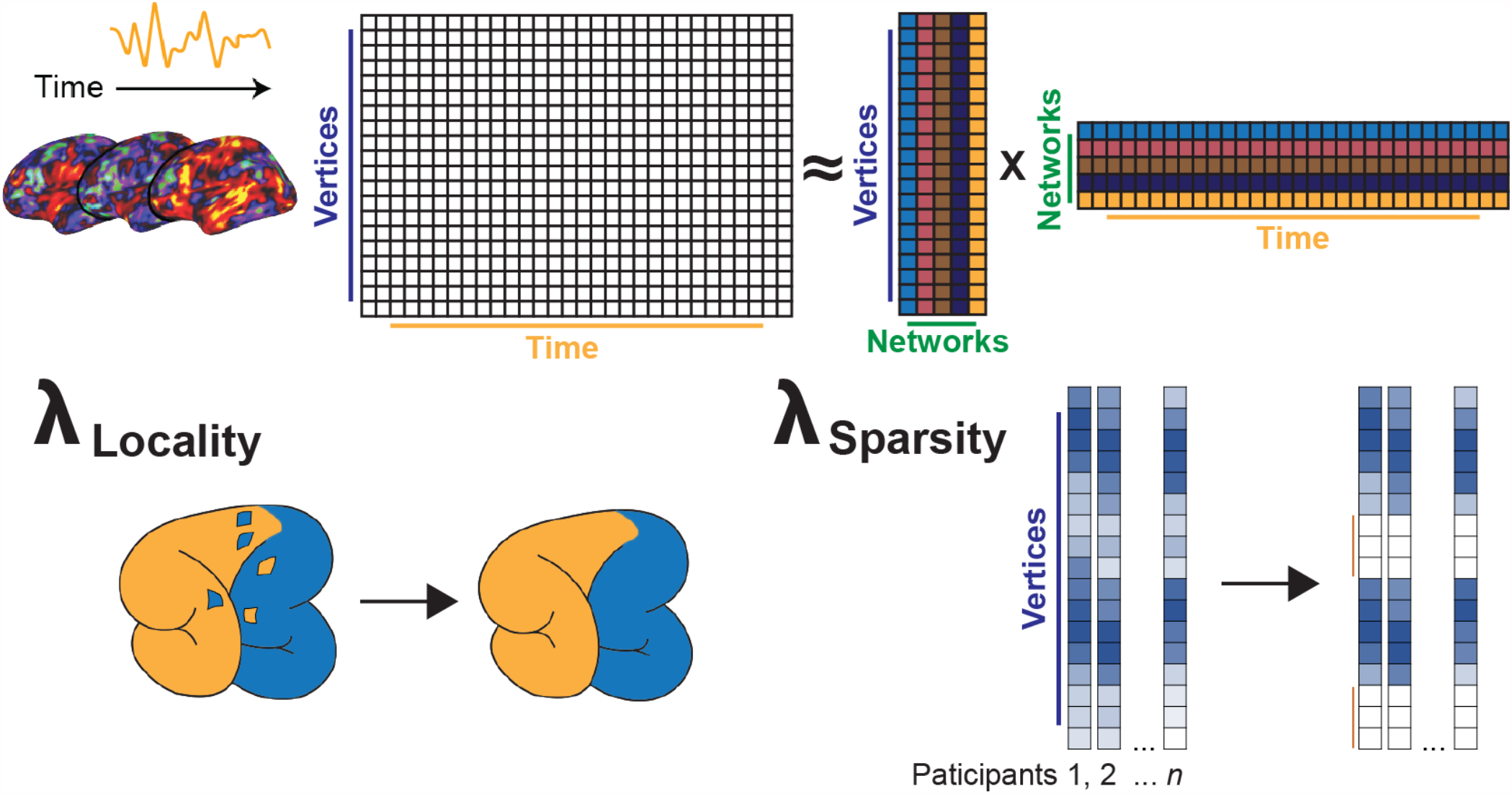
Non-negative matrix factorization (NMF) for functional brain networks. Overview of the NMF procedure. NMF leverages non-negative data, depicted here as values at each vertex over a functional time series, and performs matrix decompositions specialized for functional brain networks. Specifically, each matrix of vertex-level values is decomposed into two matrices: one representing latent functional network loadings across vertices, and the other representing loadings across time. Primarily, the cost function of NMF is reconstruction error: functional network distributions that minimize reconstruction error are preferred. In addition to reconstruction error, penalty terms encouraging spatial locality (λ_Locality_) and groupwise spatial sparsity (λ_Sparsity_) are enforced.

**Figure S2:**
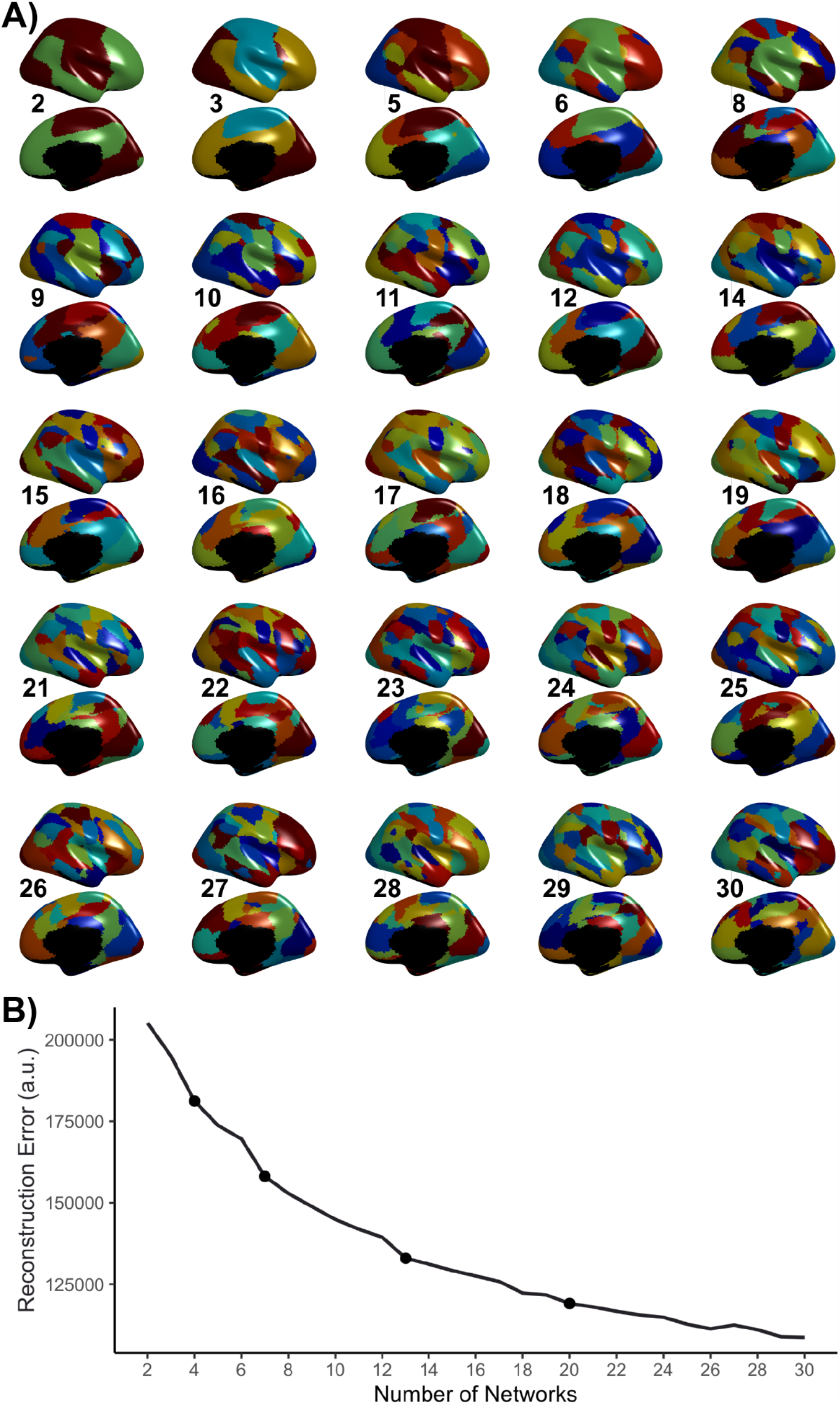
Group consensus functional network atlases. **A)** Group consensus atlases for all scales not depicted in the main text. **B)** Reconstruction error associated with each topological scale, averaged across participants. Reconstruction error descends smoothly from K=2 to K=30, suggesting that no single scale predominantly captures functional network organization. Scales chosen for visualization in the main text are demarcated with circles.

**Figure S3:**
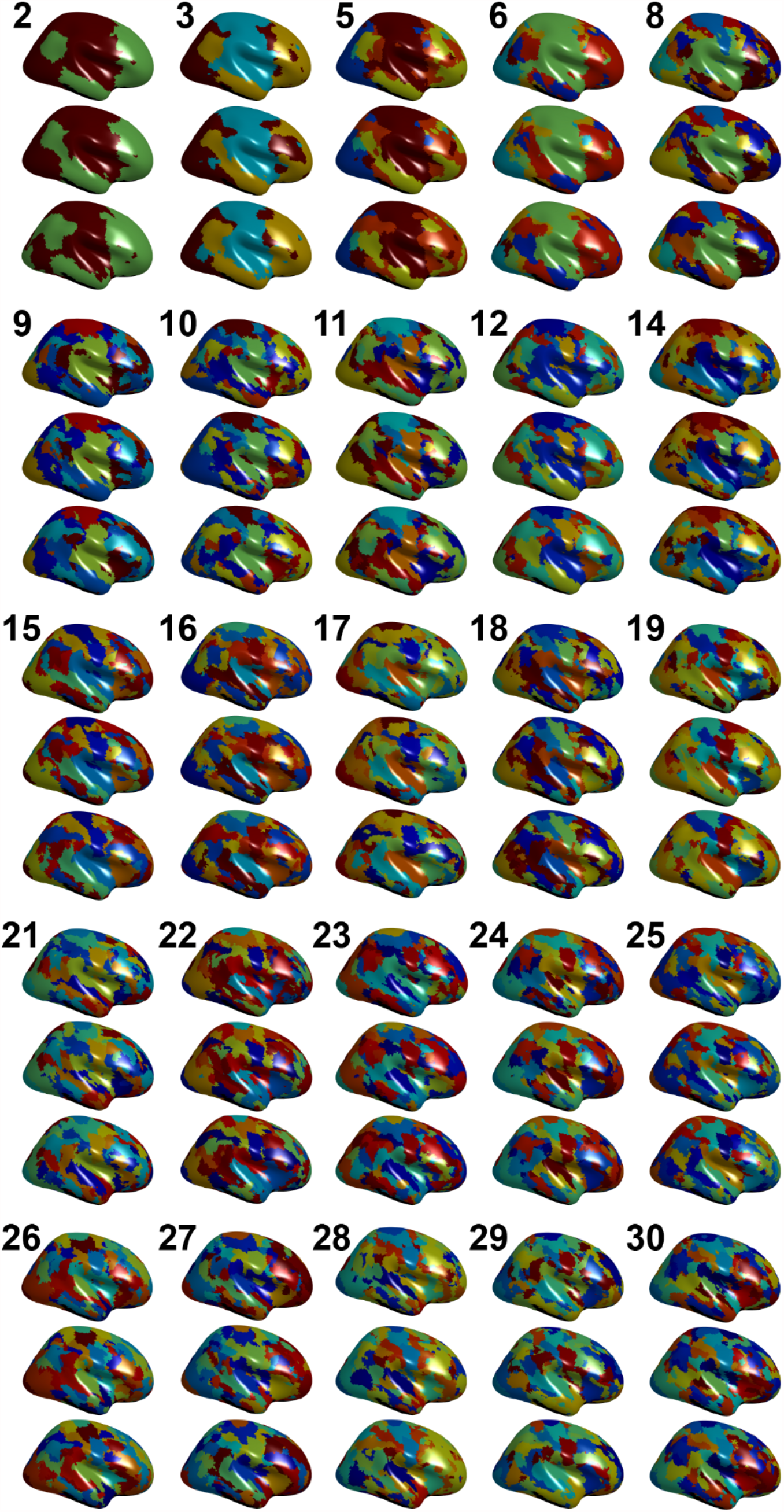
Exemplar personalized functional networks over scales. Personalized functional networks for all scales not displayed in the main text; the same example individuals from the main text are depicted in the same order.

**Figure S4:**
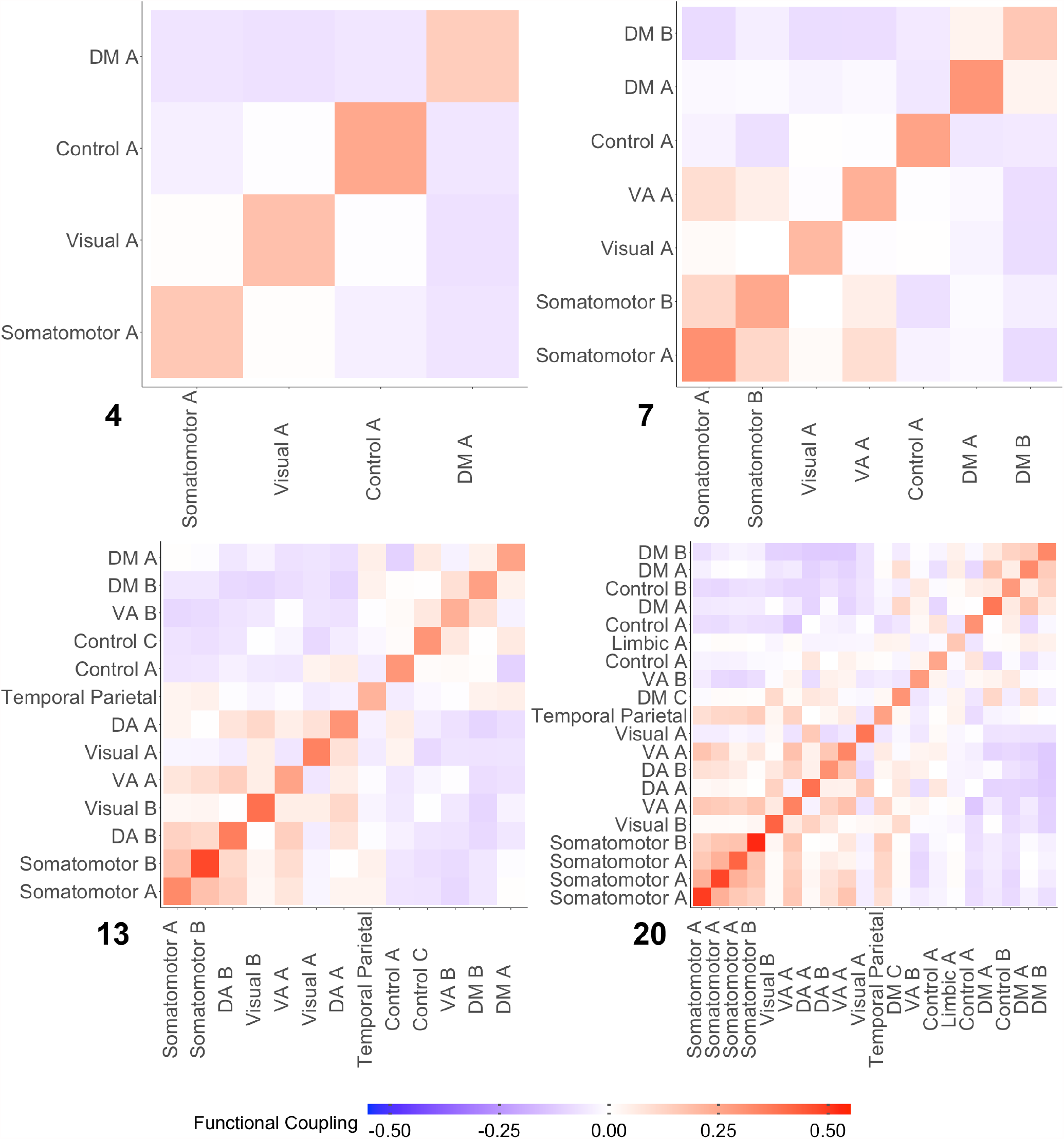
Average functional coupling of personalized networks over example scales. Functional connectivity matrices at the network-level at each scale. Network labels are derived from the maximal spatial overlap exhibited by each network with the 17-network solution from Yeo et al., 2011.

**Figure S5:**
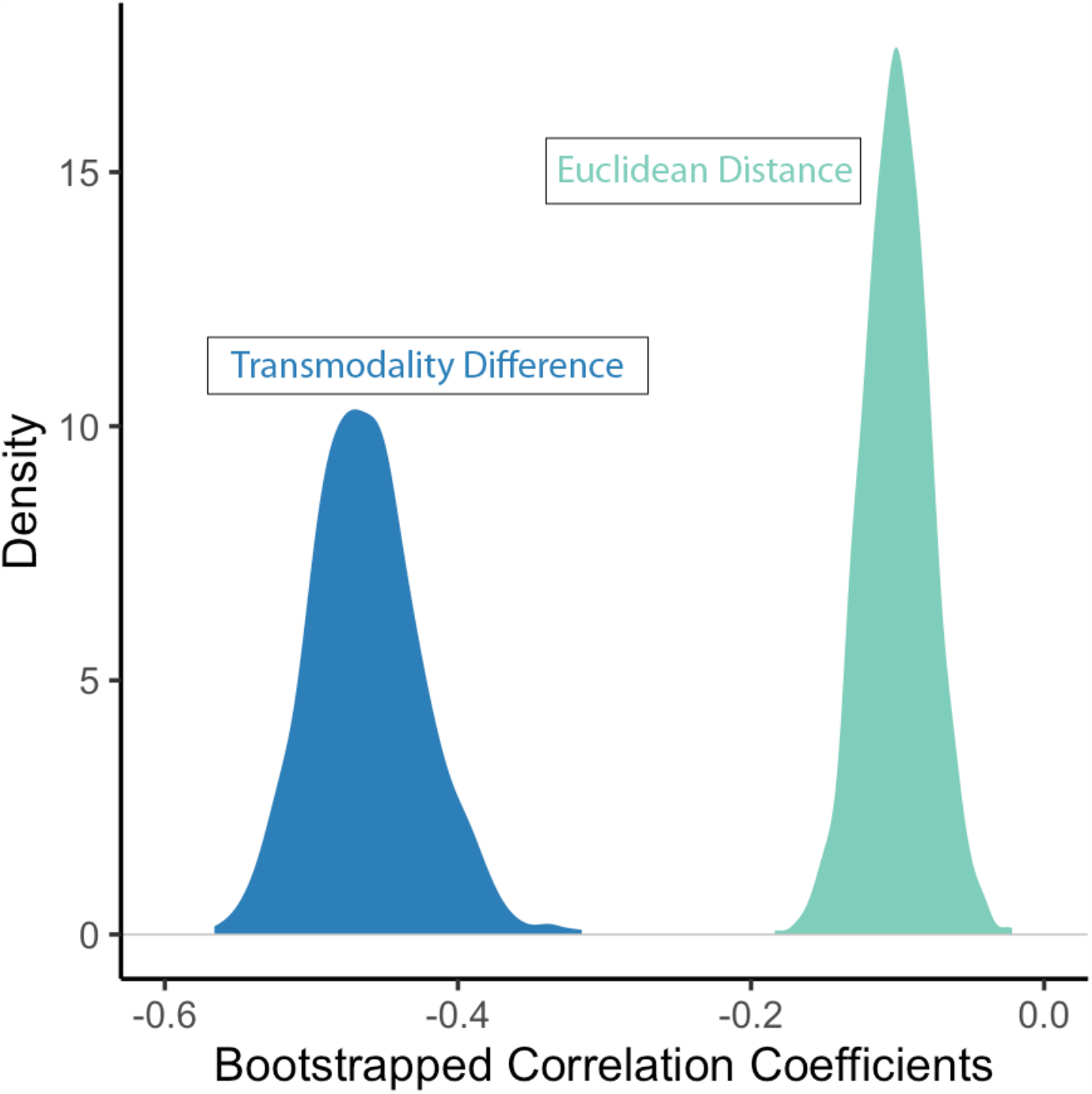
Transmodality difference and Euclidean distance between networks relates to developmental changes in between-network coupling. Functional network edge development co-varied with difference in transmodality values between networks (blue) as well as the physical distance between networks (teal). Euclidean distance is significantly negatively related to observed developmental effects, such that networks located near to each other tend to have increased coupling with age, more distant networks tend to decouple. However, these distant-dependent effects are much weaker than the effect of each network’s relative position on the sensorimotor to association axis, with transmodality difference being more strongly related to age effects across all bootstrap resamples.

## METHOD DETAILS

### Participants

A total of 1,601 participants were studied as part of the Philadelphia Neurodevelopmental Cohort (Satterthwaite et al., 2014). We excluded 340 participants due to treatment with psychoactive medications, prior inpatient psychiatric treatment, or incidentally encountered structural brain abnormalities. Among the 1,261 participants eligible for inclusion, 54 more were excluded from analyses due to low quality T1-weighted images or low quality FreeSurfer reconstructions. Of the 1,207 subjects with useable T1 images and adequate FreeSurfer reconstructions, 514 more participants were excluded for missing functional data or poor functional image quality. For inclusion in analyses, all participants were required to have three functional runs that passed quality assurance. As prior, (Ciric et al., 2018; Satterthwaite et al., 2013a), a functional run was excluded if mean relative root mean square (RMS) framewise displacement was higher than 0.2mm, or it had more than 20 frames with motion exceeding 0.25mm. This set of exclusion criteria resulted in a final sample of 693 participants with a mean age of 15.93 years (*SD* = 2.33); the sample included 301 males and 392 females. All subjects or their parent/guardian provided informed consent, and minors provided assent. All study procedures were approved by the institutional Review Boards of both the University of Pennsylvania and the Children’s Hospital of Philadelphia.

### Image acquisition

As previously described (Satterthwaite et al., 2014), all MRI scans were acquired using the same 3T Siemens Trim Trio whole-body scanner and 32-channel head coil at the Hospital of the University of Pennsylvania.

#### Structural MRI

Prior to functional MRI acquisitions, a 5-minute magnetization-prepared, rapid acquisition gradient-echo T1-weighted (MPRAGE) image (TR = 1810 ms; TE= 3.51 ms; TI = 1100 ms, FOV = 180 × 240 mm^2^, matrix = 192 × 256, effective voxel resolution = 0.9 × 0.9 × 1 mm^3^) was acquired.

#### Functional MRI

We used one resting-state and two task-based (*n*-back and emotion identification) fMRI scans for the current study. All fMRI scans were acquired with the same single-shot, interleaved multi-slice, gradient-echo, echo planar imaging (GE-EPI) sequence sensitive to BOLD contrast with the following parameters: TR = 3000 ms; TE = 32 ms; flip angle = 90° ; FOV = 192 × 192 mm^2^, matrix = 64 × 64; 46 slices; slice thickness/gap = 3/0 mm, effective voxel resolution = 3.0 × 3.0 × 3.0 mm^3^. Resting-state scans consisted of 124 volumes, while the *n*-back and emotion recognition scans consisted of 231 and 210 volumes, respectively. Further details regarding the *n*-back (Satterthwaite et al., 2013b) and emotion recognition (Wolf et al., 2015) tasks have been described in prior publications.

#### Field map

A B0 field map was derived for application of distortion correction procedures, using a double-echo, gradient-recalled echo (GRE) sequence: TR = 1000ms; TE1 = 2.69ms; TE2 = 5.27ms; 44 slices; slice thickness/gap = 4/0 mm; FOV = 240mm; effective voxel resolution = 3.8 × 3.8 × 4 mm.

#### Scanning procedure

Before scanning, to acclimate subjects to the MRI environment, a mock scanning session where subjects practiced the task was conducted using a decommissioned MRI scanner and head coil. Mock scanning was accompanied by acoustic recordings of the noise produced by gradients coils for each scanning pulse sequence. During these sessions, feedback regarding head movement was provided using the MoTrack motion tracking system (Psychology Software Tools). Motion feedback was given only during the mock scanning session. To further minimize motion, before data acquisition, participants’ heads were stabilized in the head coil using a single foam pad over each ear and a third over the top of the head.

### Image processing

#### Preprocessing

Structural images were processed with FreeSurfer (version 5.3) to allow for the projection of functional timeseries to the cortical surface (Fischl, 2012). Functional images were processed using a top-performing preprocessing pipeline implemented using the eXtensible Connectivity Pipeline (XCP) Engine (Ciric et al., 2018), which includes tools from FSL (Jenkinson et al., 2012; Smith et al., 2004) and AFNI (Cox, 1996). This pipeline included (1) correction for distortions induced by magnetic field inhomogeneity using FSL’s FUGUE utility, (2) removal of the initial volumes of each acquisition, (3) realignment of all volumes to a selected reference volume using FSL’s MCFLIRT, (4) interpolation of intensity outliers in each voxel’s time series using AFNI’s 3dDespike utility, (5) demeaning and removal of any linear or quadratic trends, and (6) co-registration of functional data to the high-resolution structural image using boundary-based registration (Greve & Fischl, 2009). Images were de-noised using a 36-parameter confound regression model that has been shown to minimize associations with motion artifact while retaining signals of interest in distinct sub-networks (Ciric et al., 2017). This model included the six framewise estimates of motion, the mean signal extracted from eroded white matter and cerebrospinal fluid compartments, the mean signal extracted from the entire brain, the derivatives of each of these nine parameters, and quadratic terms of each of the nine parameters and their derivatives. Both the BOLD-weighted time series and the artifactual model time series were temporally filtered using a first-order Butterworth filter with a passband between 0.01 and 0.08 Hz to avoid mismatch in the temporal domain (Hallquist et a., 2013). Furthermore, to derive time series that were more comparable across runs, the task activation model was regressed from *n*-back and emotion identification fMRI data (Fair et al., 2007b). The task activation model and nuisance matrix were regressed out using AFNI’s 3dTproject.

For each modality, the fMRI timeseries of each participant was projected to their own FreeSurfer surface reconstruction and smoothed on the surface of this reconstruction with a 6-mm full-width half-maximum kernel. The smoothed data was projected to the *fsaverage5* template, which has 10,242 vertices on each hemisphere (18,715 total vertices after removing the medial wall). Finally, we concatenated the three fMRI acquisitions, yielding a timeseries of 27 minutes and 45 seconds in total (555 volumes). As prior, we removed vertices with low signal-to-noise ratio (SNR; Gordon et al., 2016; Ojemann et al., 1997; Wig et al, 2014). We used the same SNR mask as in our prior work, which used the same dataset (Cui et al., 2020). After masking, 17,734 vertices remained for subsequent analyses.

#### Regularized non-negative matrix factorization

As previously described in detail (Li et al., 2017), we used non-negative matrix factorization (NMF; Lee and Seung, 1999) to derive personalized functional networks. The NMF method decomposes the time series by positively weighting cortical vertices that covary, leading to a highly specific and reproducible parts-based representation (Lee and Seung, 1999; Soritas et al., 2017). Our approach was enhanced by a group consensus regularization term that preserves inter-individual correspondence, as well as a data locality regularization term to mitigate imaging noise, improve spatial smoothness, and enhance functional coherence of personalized functional networks (se Li et al., 2017 for details of the method; see also: https://github.com/hmlicas/Collaborative_Brain_Decomposition). As NMF requires non-negative input, we shifted the timeseries of each vertex linearly to ensure all values were positive. Finally, all vertex timeseries were normalized to their maximum values such that all values ranged between 0 and 1.

Given a group of *n* subjects, each having fMRI data X^*i*^ ∈ R^×^, *i* = 1, …, *n*, consisting of *S* vertices and *T* time points, we aimed to find *K* non-negative functional networks *V*^*i*^ = (*V*^*i*^ _*s,k*_)∈*R*^*S*×*K*^ and their corresponding time courses *U*^*i*^ = (*U*^*i*^ _*t,k*_)∈*R*^*T*×*K*^ for each subject, such that

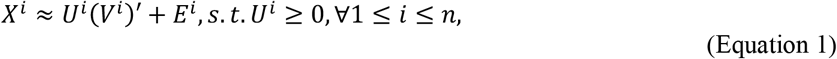

Where (*V*^*i*^)^′^ is the transpose of (*V*^*i*^) and *E*^*i*^ is independently and identically distributed residual noise following a gaussian distribution. Both *U*^*i*^ and *V*^*i*^ were constrained to be non-negative so that each functional network did not contain anti-correlated functional units. A group consensus regularization term was applied to ensure inter-individual correspondence, which was implemented as a group-sparsity term on each column of *V*^*i*^ and formulated as

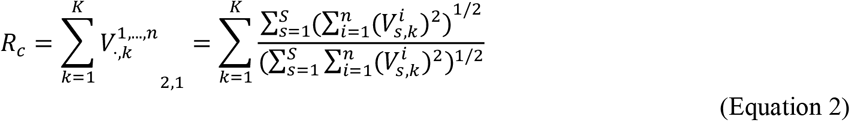

The data locality regularization term was applied to encourage spatial smoothness and coherence of the functional networks using graph regularization techniques (Cai et al., 2011). The data locality regularization term was formulated as

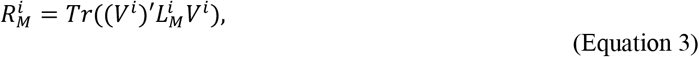

where 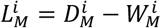 is a Laplacian matrix for subject *i*, 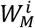 is a pairwise affinity matrix to measure spatial closeness or functional similarity between different vertices, and 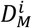 is its corresponding degree matrix. The affinity between each pair of spatially connected vertices (here, vertices *a* and *b*) was calculated as 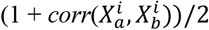, where 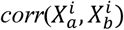 is the Pearson correlation coefficient between time series 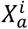 and 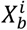; the pairwise affinity between non-connected vertices was set to zero so that 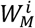 would be sparse. We identified personalized functional networks by optimizing a joint model with integrated data fitting and regularization terms formulated as

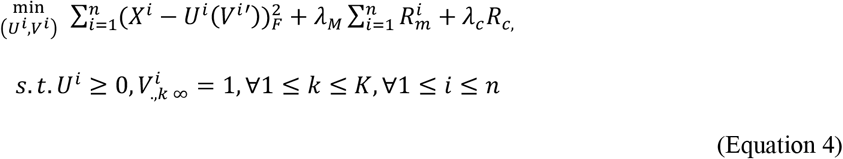

where *λ*_*M*_ = *β* × (*T* /*K* × *n*_*m*_) and *λ*_*c*_ = *α* · (*n* · *T* /*K*) are used to balance the data fitting, data locality, and group consensus regularization terms, *n*_*m*_ is the number of neighboring vertices, and *α* and *β* are free parameters. For this study, we used previously validated parameters (*α, β* = 1,10 for group consensus partitions, *α, β* = 10,1 for individualized partitions; Li et al., 2017; Cui et al., 2020) across 29 values of *K* (*K=*2 to *K*=30) corresponding to 29 scales of cortical organization.

#### Defining personalized networks

Our approach for defining personalized networks included three steps. In the first two steps, a group consensus atlas was created. In the third step, this group atlas was used to initialize network personalization for each participant at each scale. Although individuals exhibit distinct network topography, broad consistencies exist from individual-to-individual (Gordon et al., 2017b, Gratton et al., 2018). By first generating a group atlas for personalization initialization, we ensured spatial correspondence across all subjects and scales. This strategy has also been applied in other studies of personalized networks (Kong et al., 2019; Wang et al., 2015). For computational efficiency and to avoid outlier-driven group atlases, a bootstrap strategy was utilized to perform the group-level decomposition multiple times on a subset of randomly selected participants. Subsequently, the resulting decompositions were fused to obtain one robust initialization. As prior (Li et al., 2017; Cui et al., 2020), we randomly selected 100 subjects and temporally concatenated their timeseries, resulting in a timeseries matrix with 55,500 rows (time-points) and 17,734 columns (vertices). We applied the above-mentioned regularized non-negative matrix factorization method with a random initialization to decompose this group-level matrix (Lee and Seung, 1999). A group-level network loading matrix *V* was acquired, which had *K* rows and 17,734 columns. Each row of this matrix represents a functional network, while each column represents the loadings of a given cortical vertex. As prior, (Li et al., 2017; Cui et al., 2020), this procedure was repeated 50 times, each time with a different subset of subjects. Accordingly, this process yielded 50 different group atlas estimations for each value of *K*.

Next, we combined the 50 group network atlases to obtain one robust group network atlas with spectral clustering at each value of *K*. Specifically, we concatenated the 50 group parcellations together across networks to obtain a matrix with 50 × *K* rows (functional networks) and 17,734 columns (vertices). Next, we calculated inter-network similarity as

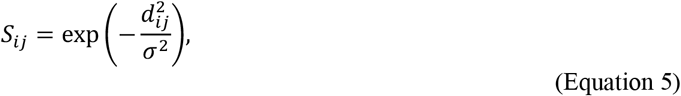

where *d*_*ij*_ = 1 – *corr*(*Network*_*i*_, *Network*_*j*_), *corr*(*Network*_*i*_, *Network*_*j*_) is a Pearson correlation coefficient between Network_i_ and Network_j_, and *σ* is the median of *d*_*ij*_ across all possible pairs of functional networks. Then, we applied normalized-cut-based spectral clustering (Cai et al., 2011) to group the 50 × *K* functional networks into *K* clusters. For each cluster, the functional network with the highest overall similarity with all other networks in the same cluster was selected as the most representative. The final group network atlas was composed of these maximally representative network estimations at each of the 29 resolutions studied.

In the final step, we derived each individual’s specific network atlas using NMF, initializing each participant-specific solution on the group consensus atlas for any given scale and optimizing NMF in accordance with each individual’s specific fMRI time series (a 555 × 17,734 matrix). See Li et al., (2017) for further optimization detail. This procedure yielded loading matrix *V*_*i*_ (*K* × 17,734 matrix) for each participant, where each row is a functional network, each column is a vertex, and the value in each cell quantifies the extent to which each vertex belongs to each network. This probabilistic (soft) definition was converted into discrete (hard) network definitions for display and calculation of network statistics by labeling each vertex in accordance with its highest loading. This procedure was repeated for all 29 network resolutions.

## QUANTIFICATION AND STATISTICAL ANALYSIS

### Calculation of variability and spatial alignments of personalized networks

In order to quantify cross-subject spatial variability in personalized networks, we calculated the median absolute deviation (MAD) of personal network loadings at each vertex across participants. MAD is a non-parametric measure of variance that does not assume a normal distribution. First, we calculated MAD for each network at each scale. Next, MAD was averaged across *K* networks to obtain a single value of MAD at each vertex for any given scale *K*.

### Principal gradient

In order to quantify cortices in terms of their position within a functional hierarchy, we used a widely adopted principal gradient of functional connectivity (Marguiles et al., 2016; https://github.com/NeuroanatomyAndConnectivity/gradient_analysis). The principal gradient is derived from the primary component of variance in patterns of whole-brain functional connectivity, and it reflects a unimodal-to-transmodal continuum of cortical function. As such, at each cortical vertex, the value of this gradient reflects the loading of that vertex onto the unimodal-to-transmodal continuum, with higher principal gradient values corresponding to higher transmodality. This map was transformed to *fsaverage5* space using *metric-resample* from Connectome Workbench. Transmodality values for each network were quantified as the average principal gradient value of each vertex within each network in group-consensus space. These network-wise transmodality values were used to analyze the spatial distribution of the effects of age and executive function, as described below.

### Reference networks

To allow for comparison with previously estimated cortical systems, we quantified the overlap of each group-consensus network with a commonly used 7 and 17-functional network parcellation (Yeo et al., 2011). To illustrate this overlap, we assigned colors to group and individualized networks in accordance with their maximum overlap with networks from the 7 and 17-network parcellations.

### Spatial permutation testing (spin test)

In order to evaluate the significance of the localization of between-participant variability (MAD) to transmodal cortical areas, we used a spatial permutation procedure called the spin test (Alexander-Bloch et al., 2018; Gordon et al., 2016; Sotiras et al., 2017; Vandekar et al., 2015; https://github.com/spin-test/spin-test). The spin test is a spatial permutation method based on angular permutations of spherical projections at the cortical surface. Critically, the spin test preserves the spatial covariance structure of the data, providing a more conservative and realistic null distribution than randomly shuffling locations. Due to varying spatial covariance structure across scales, we conducted separate spin tests at each scale.

### Modeling the association of scale with MAD-principal gradient co-localization

To account for potential non-independence of MAD-principal gradient correlations across scales, significance testing was performed using non-parametric bootstrap resampling. Specifically, we re-calculated MAD and the subsequent spatial correlation with the principal gradient at each scale across 1,000 bootstrap resamples to generate a bootstrapped confidence interval of the second-order relationship between network scale and the MAD-principal gradient correlations.

### Quantification of between-network coupling

We used functional connectivity (FC) to quantify inter-regional coupling in processed BOLD signal.

Specifically, we calculated between-network FC at three levels of analysis: network, edge, and vertex. At all levels, FC was quantified as the Pearson correlation between BOLD timeseries. At the network level, between-network connectivity was quantified as a network’s mean correlation with all other networks. At the edge level, between-network connectivity was quantified as the mean vertex-by-vertex correlation between vertices in both networks. At the vertex-level, we evaluated each vertex’s average correlation to vertices from all other networks. Between-network coupling at each level was quantified separately at each scale for each participant.

### Developmental analyses

#### Developmental modeling

Developmental effects were estimated using generalized additive models (GAMs; Wood, 2001, 2004) with penalized splines in R (Version 3.6.3) using the *mgcv* package (R Core Team, 2013; Wood, 2011). To avoid over-fitting, nonlinearity was penalized using restricted maximum likelihood (REML). Participant sex and in-scanner head motion were included as covariates within each GAM. Head motion was quantified as the mean framewise root-mean-square displacement across the three functional runs for each subject. Age was modeled using a penalized thin-plate regression spline; covariates were modeled as parametric regressors. This model can summarized using the formula in equation 6:

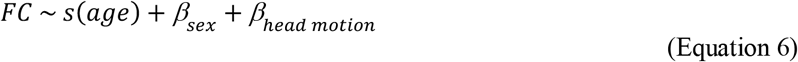

To quantify the effect sizes of each age spline, we calculated the change in adjusted *R*^2^(Δ *R*^2^_*adj*._) between the full model and a nested model that did not include an effect of age. Statistical significance was assessed using analysis of variance (ANOVA) to compare the full and nested models. Because Δ *R*^2^_*adj*._. describes effect size but not direction (i.e., increasing or decreasing FC with age), we extracted and applied the sign of the age coefficient from an equivalent linear model as in prior work (Cui et al., 2020). To estimate windows of significant age-related change for each network-level model, we calculated the age range for which the 95% confidence interval of estimated age splines did not include 0 (Larsen et al., 2020; Pines et al., 2020). To calculate the intervals, we used the *gratia* package in R (Simpson, 2018). Multiple comparisons were controlled for with the false discovery rate (FDR) correction (*q* < .05).

#### Modeling the distribution of developmental effects across the principal gradient

After analyzing the effect of age on between-network FC, we sought to evaluate the spatial distribution of age effects along the principal gradient. At the network level, we extracted the mean transmodality value for each network at each scale and regressed these values on the corresponding pattern of age effects (Equation 7).

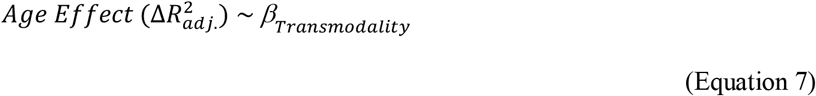

To account for potential non-independence of age effects across scales, significance testing was performed using non-parametric bootstrap resampling. Specifically, we re-calculated the age effects for each network and the resulting transmodality relationship across 1,000 bootstrap resamples to generate a bootstrapped confidence interval. The effect size of the second-order model was also described as a Spearman’s correlation coefficient.

We next evaluated how the magnitude of the age effects corresponded to the span of each edge (between-network connection) across the transmodality gradient. We modeled this effect in two ways. First, we calculated the difference in the transmodality gradient values for each pair of networks at each scale (“transmodality difference”) and regressed this difference on the age effects from the edge-wise developmental models (Equation 8).

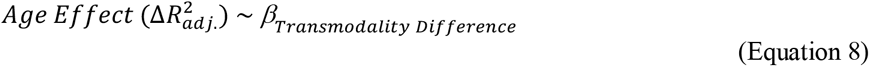

As above, significance was evaluated using non-parametric bootstrap resampling. As a sensitivity analysis, we repeated this procedure using the average Euclidean distance between vertices in the two networks comprising each edge. Second, we sought to visualize the interaction between transmodality difference and age-related changes in coupling across network edges spanning different portions of the principal gradient. In order to continuously model the relation between age-related changes in coupling and transmodality difference across the principal gradient, we fit a bivariate smooth interaction. Specifically, we modeled the effect of transmodality on the edge-level age effects using a tensor product smooth (Wood, 2006) as in equation 9.

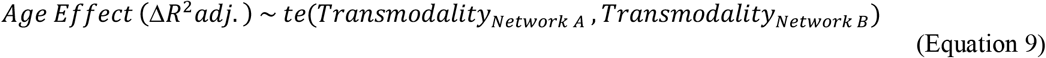

To verify the statistical significance of this model, we performed the same non-parametric bootstrap procedure as above using a simplified linear interaction model.

#### Modeling scale-dependent developmental effects

In order to quantify and localize the scale-dependence of developmental changes in between-network coupling, we modeled the impact of scale on coupling at each vertex. Model formulas and initial model fits were estimated using GAMs (Equation 10).

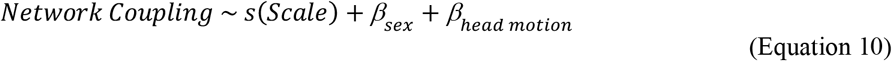

GAM-derived coefficient estimates for scale, sex, and head motion were used to initialize generalized estimating equations (GEEs). GEEs enabled us to account for the covariance between same-subject measurements across scales without assuming independence of these observations. At each vertex, the effect of scale was assessed for statistical significance via a joint Wald test that compared the full model to a nested model that did not include an effect of scale.

Age-by-scale interactions were modeled using the same procedure. First, GAMs were used to generate initial model fits. Age-by-scale interactions were modeled as a bivariate tensor product interaction (*ti* in mgcv) as in equation 11.

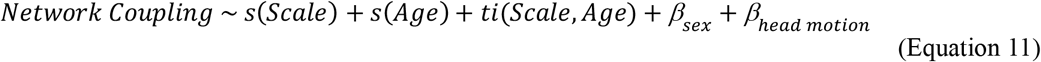

Again, GEEs were used to account for the covariance between same-subject measurements across scales without assuming independence. Statistical significance was evaluated with a joint Wald test that compared the full model to a nested model that did not include a bivariate interaction term.

Finally, to further understand scale-dependent age effects within areas exhibiting age-by-scale interactions, we compared network level developmental effects across scales for networks that fall at opposite ends of the principal axis. We grouped networks by their maximum overlap with the higher-resolution reference atlas (the 17 network solution provided by Yeo et al.) and calculated average transmodality values for each group of reference networks. The lowest (Somatomotor-A) and highest (Default mode-B) transmodality networks were chosen to depict differential scale-dependence across the principal gradient. To illustrate the effect of scale, we fit a penalized spline to the relationship between scale and observed age effects for each network within each group.

### Analyses of executive function

#### Cognitive assessment

The Penn computerized neurocognitive battery (Penn CNB) was administered to all participants as part of a session separate from neuroimaging. The CNB consists of 14 tests adapted from tasks applied in functional neuroimaging to evaluate a broad range of cognitive domains (Gur et al., 2012). These domains include executive function (abstraction and mental flexibility, attention, working memory), episodic memory (verbal, facial, spatial), complex cognition (verbal reasoning, nonverbal reasoning, spatial processing), social cognition (emotion identification, emotion differentiation, age differentiation), and sensorimotor speed. Accuracy for each test was *z*-transformed. As previously described in detail, factor analysis was used to summarize these accuracy scores into three factors (Moore et al., 2015), including executive and complex cognition, episodic memory, and social cognition. Here, we focused on the executive and complex cognition accuracy factor score.

#### Cognitive modeling

Analyses of associations with cognition were executed using GAMs, as described above for developmental analyses. Specifically, EF was modeled using a penalized regression spline, while covarying for age using a penalized regression spline; participant sex and mean head motion were included as linear covariates (Equation 12).

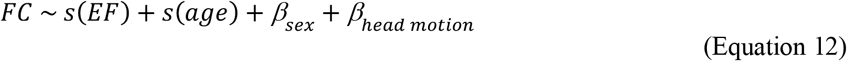

As for developmental analyses, we calculated the effect size as the change in adjusted *R*^2^ between the full model and a nested model that did not include the effect of EF (Δ *R*^2^_*adj*._).

#### Linking associations with EF to the principal gradient of brain organization

After analyzing the effect of cognition on between-network FC, we sought to evaluate the distribution of EF effects across the sensorimotor to association axis. At the network level, we extracted the mean transmodality value for each network at each scale and compared these values to the corresponding pattern of associations between between-network coupling and EF. As for previous developmental analyses, in order to assess the statistical significance of EF effect-transmodality correspondence, we also evaluated a second-order model over 1,000 bootstrap resamples. However, here we also included quadratic term (Equation 13).

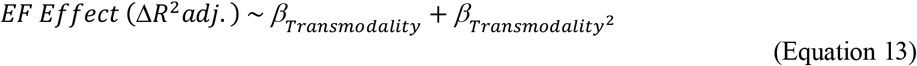

The resulting bootstrapped confidence intervals for *β*_*Transmodality*_ and 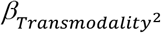 were then used for significance testing of these second-order effects.

#### Modeling scale-dependent cognitive effects

In order to quantify and localize the scale-dependence of associations between EF and between-network coupling, we modeled the impact of scale at each vertex. EF-by-scale interactions were modeled using the same procedure as for developmental models. First, GAMs were used to generate initial model fits. EF-by-scale interactions were modeled as a bivariate tensor product interaction (*ti* in mgcv) as in equation 14.

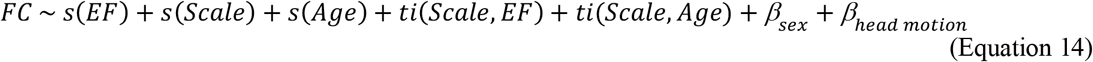

Again, GEEs were used to account for the covariance between same-subject measurements across scales without assuming independence. Statistical significance was evaluated with a joint Wald test that compared the full model to a nested model that did not include a bivariate interaction term.

Finally, to further understand scale-dependent cognitive effects within areas exhibiting EF-by-scale interactions, we compared network level cognitive effects across scales for networks that fall at opposite ends of the principal axis. To model the effect of scale, we fit a penalized spline to the relationship between scale and observed cognition effects for the lowest (Somatomotor-A) and highest (Default mode-B) transmodality networks.

#### Multivariate EF predictions

As a final step, we sought to assess the degree to which multivariate patterns of functional edge coupling across scales jointly explains individual differences in executive function. To do this, we used ridge regression. We iteratively fit a regression model to two thirds of our sample (462 participants) and predicted executive function scores from functional coupling data in the left-out testing third of participants (231 participants). In each iteration, we used nested parameter optimization. Specifically, coefficients for each edge were fit with the 1^st^ third of the sample and then the L2 penalty term was selected based on predictions in the 2^nd^ third of the sample. Finally, the degree to which functional coupling explains EF was calculated using the unseen 3^rd^ third of the sample. In that left-out data that was not used in model training, we calculated the correlation between actual and predicted EF, as well as the mean squared error (MSE). We repeated this process 100 times to ensure that specific sample splits were not driving results, and averaged predictions across iterations. To evaluate statistical significance of these predictions, we used permutation testing. Specifically, we repeated this process 1000 times, and compared our outcome measure (correlation of predicted vs. actual EF) versus a distribution of models where EF scores had been permuted across participants.

### Mediation analyses

To test mediation effects, we applied a product-of-paths mediation framework where developmental changes in between-network coupling mediated age-related increases in EF (see Figure 7B). Specifically, for each network, we posited that the effect of age on EF (*c* pathway) was mediated by the effect of age on between-network coupling (*a* pathway) and the effect of between-network coupling on EF (*b* pathway). All mediation pathways were estimated with linear models in the *lavaan* package in R. We estimated the mediation effect for each network at each scale as the product of the *a* and *b* pathways (*a* pathway ^*^ *b* pathway). Sex and in-scanner motion were included as covariates in all mediation models.

Finally, in order to evaluate the relationship between the principal gradient and observed mediation effects, we regressed network transmodality values on the AB path coefficients across networks and scales using equation 15.

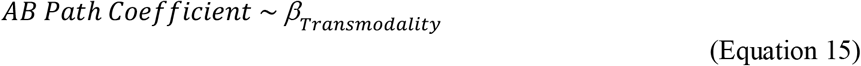

As prior, statistical significance was evaluated using non-parametric bootstrap resampling (1,000 iterations).

## DATA AND CODE AVAILABILITY

The PNC data is publicly available in the Database of Genotypes and Phenotypes: accession number: phs00607.v3.p2; https://www.ncbi.nlm.nih.gov/projects/gap/cgi-bin/study.cgi?study_id=phs000607.v3.p2. All analysis code is available here https://github.com/PennLINC/multiscale, with detailed explanation provided at https://pennlinc.github.io/multiscale/.

